# geneRFinder: gene finding in distinct metagenomic data complexities

**DOI:** 10.1101/2020.08.21.262147

**Authors:** Raíssa Silva, Kleber Padovani, Fabiana Góes, Ronnie Alves

## Abstract

**Motivation:** Microbes perform a fundamental economic, social and environmental role in our society. Metagenomics makes it possible to investigate microbes in their natural environments (the complex communities) and their interactions. The way they act is usually estimated by looking at the functions they play in those environments and their responsibility is measured by their genes. The advances of next-generation sequencing technology have facilitated metagenomics research however it also create a heavy computational burden. Large and complex biological datasets are available as never before. There are many gene predictors available which can aid gene annotation process though they lack of handling appropriately metagenomic data complexities. There is no standard metagenomic benchmark data for gene prediction. Thus, gene predictors may inflate their results by obfuscating low false discovery rates.

**Results:** We introduce geneRFinder, a ML-based gene predictor able to outperform state-of-the-art gene prediction tools across this benchmark by using only one pre-trained Random Forest model. Average prediction rates of geneRFinder differed in percentage terms by 54% and 64%, respectively, against Prodigal and FragGeneScan while handling high complexity metagenomes. The specificity rate of geneRFinder had the largest distance against FragGeneScan, 79 percentage points, and 66 more than Prodigal. According to McNemar’s test, all percentual differences between predictors performances are statistically significant for all datasets with a 99% confidence interval.

**Conclusions:** We provide geneRFinder, a approach for gene prediction in distinct metagenomic complexities, available at github.com/railorena/geneRFinder, and also we provide a novel, comprehensive benchmark data for gene prediction — which is based on The Critical Assessment of Metagenome Interpretation (CAMI) challenge, and contains labeled data from gene regions – avaliable at sourceforge.net/p/generfinder-benchmark.

## Background

Prokaryotic organisms are found everywhere, in soil, water, animals, being responsible for key roles in their survival and maintenance. Bacteria in the intestines of humans, for example, not only aid in the digestion of food but also greatly interfere with the vital systems of human beings, such as the immune system [1], thus making humans highly dependent on a perfect balance among microorganisms interaction.

Identifying which prokaryotes coexist in environments and in-depth knowledge about these microorganisms enables valuable scientific discoveries that can benefit all ecosystems related to these microorganisms, especially on humans by advances in areas of disease prevention and cure [2]. Describing genes in prokaryotes genomes is one way to understand how these microorganisms play in complex systems.

This identification, also referred to as annotation, is commonly performed with the aid of prediction systems that locate genes along genomes using a reference database composed of genes previously annotated in related genomes. Although gene annotation has grown in recent years, there are still countless genes that have not been annotated, thus making predictions solely based on available known reference genomes quite limited and will not always be sufficient to describe the main role of these microorganisms.

Gene prediction based on the structures of the analyzed genomic sequences - also known as ab initio [3] - is a way to identify genes independently and more aligned with the current reality of prokaryotic genomic studies - which, in turn, estimates, it has information on only about 1% of existing species [4].

Ab initio prediction is commonly based on the identification of protein-coding sequences (CDS) contained in genes and can be performed by the Open Read Frame (ORF) extraction method [5]. The term ORF corresponds to a portion of the genome - that is, a genomic sequence - initiated and terminated by a specific combination of nucleotides, known as start and stop codon respectively. However, the prediction process is not so trivial because not every ORF found in the genome corresponds to a CDS [6]. Thus, ORF extraction alone does not satisfy the sufficient condition for CDS identification, requiring that other sequence properties need to be considered for gene prediction.

Although there are well-used and well-performing tools for gene prediction, such as FragGeneScan [7] and Prodigal [8], this task is still a challenge. This difficulty becomes greater when gene prediction must be performed in environmental metagenomic samples. As an example, soil samples present a wide diversity of species linked to distinct metagenomics complexities [9].

Metagenomic samples with a high number of species are commonly referred to as high complexity samples and therefore contain high genomic diversity. Using traditional metagenomic data analysis procedures, this diversity can produce inconsistencies [10] - due to the mixing of genetic information - impacting on the quality of gene prediction tools.

Metagenomic data complexity is a topic superficially considered in the evaluation of gene predictors, possibly justifying by the lack of metagenomic dataset benchmarks for such use. This scenario exposes an interesting gap in the effectiveness of the performance analysis of these tools and highlights the need to create fair benchmarks for this purpose.

The inability to characterize non-coding sequences, or intergenic region, remains another challenge. Different that was previously believed, non-coding regions - where it is possible to find sequences as translation initiation site, promoters and terminators [11] - have important information capable of distinguishing the pathogenic and non-pathogenic strains [12], as well as other functions, however, our knowledge about the exact biological functions of these sequences is limited [13] and needs further investigation.

In this paper, we propose geneRFinder, an ab initio gene prediction tool capable of identifying CDS and intergenic region within sequences distinct metagenomic complexities. This tool was built on the Random Forest classifier model due to its good performance when compared to other known classification methods applied to the discovery of genes in metagenomic data [14]. Additionally, we produced and provided a metagenomic gene prediction benchmark for validation of gene prediction tools, that is composed by 9 datasets - 4 manually produced datasets and 5 datasets derived from the benchmark data provided by the first edition of the well-known Critical Assessment of Metagenome Interpretation (CAMI) challenge.

## Implementation

The geneRFinder is an ORF extraction based tool capable of identifying coding sequences and intergenic regions in metagenomic sequences, being able to predict independently based on signal capture from those regions. As it will be presented in more detail in the following subsections, properties of sequences are extracted from ORFs that are then transformed into numerical vectors to be learned by a Random Forest model [15]. Such model was trained and validated in datasets of microorganisms that had complete genome and annotations. The final model was tested on independent datasets having different genome complexities and sequences sizes.

### Training and Validation Datasets

Complete genomes and their complementary information provided by the NCBI [16] genome repository were used, including annotated CDS and the gene and CDS mapping table for each organism, to create training and validation datasets. ORFs located in the genomes were extracted and, for each of them, were assigned the corresponding label - positive for CDS (and internal ORFs) and negative for not being a coding sequence, according to the respective NCBI mapping table, thus, was recognized as a non-coding sequence everything that is between CDS, for example, translation initiation site. The ORF extraction process considered as ORF the sequences found in the genomic sequences that had ATG as start codon and TAG, TGA or TAA for the stop codon.

Initially, 20 complete genomes were used to tune parameters that contributed to the differentiation of gene and intergenic region, as well as to identify the characteristics of sequences useful to generate the learning model. This model was then validated on 5 different genomes of the training set introduced in [17]. Next, a more enriched model was built, consisting of 129 complete genomes and their respective annotations, of which 11 are archaeas and 118 are bacteria. From these genomes, 712,868 sequences were extracted, 356,443 of which correspond to CDS, hereinafter referred to as positive instances, and 356,425 to intergenic regions (negative instances). The genomes names, the taxonomy ID and the taxonomy level are depicted in Table 1, 2 and 3 of the supplementary material.

### Test Datasets

The test dataset was built using 12 public genomes and respective annotations, 3 archaeas and 9 bacteria, following the same methodology described in the previous section to obtain the ground truth. From these organisms, 31,507 positive and 23,473 negative ORFs were extracted, totaling 54,980 sequences to be predicted. In order to make a fair comparison performance analysis with the currently state-of-the-art gene prediction tools - namely FragGeneScan, Orphelia [18], MetaGene [19] and Prodigal - the 12 most frequently used genomes listed in their respective publications were selected for further analysis (Figure 1).

**Figure 1.**
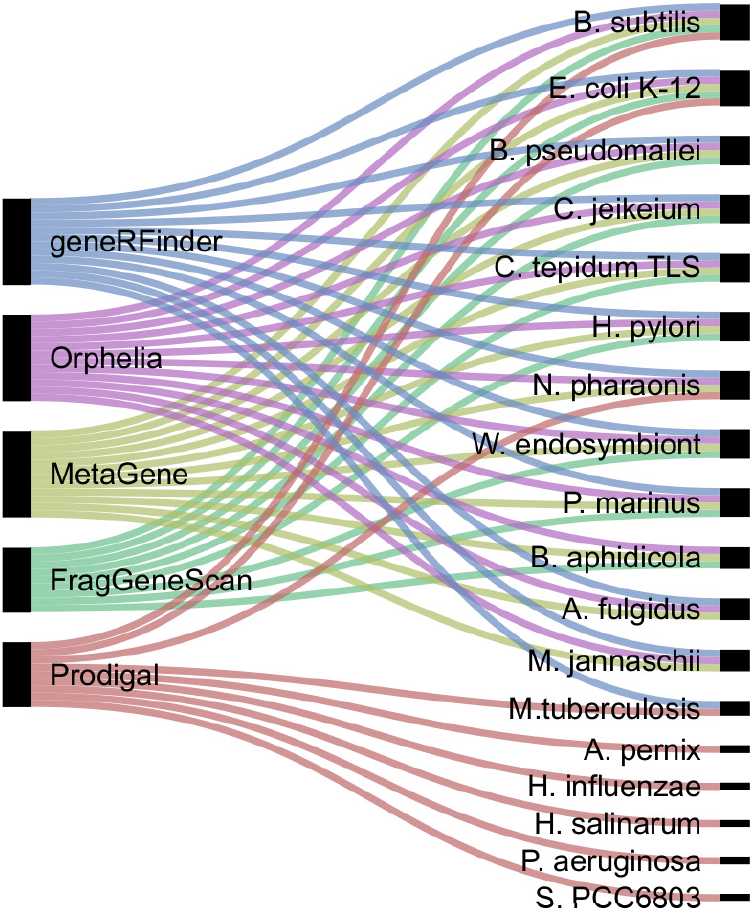
The set of genomes frequently used by gene prediction tools. On the left gene predictors and on the right genomes.

#### Benchmark dataset from CAMI

geneRFinder was also tested on datasets extracted from CAMI [15], a metagenomic benchmark that features datasets for assembly and binning evaluation of samples in three distinct complexities (low, medium and high), containing sequences of bacteria, archaea and viruses. The benchmark introduces three assemblies - each for a level of complexity - considered optimal. Some information about assemblies is provided in Table 1. The values of N50, L50 and contig numbers were analyzed by the Metaquast tool [20].

**Table 1.**
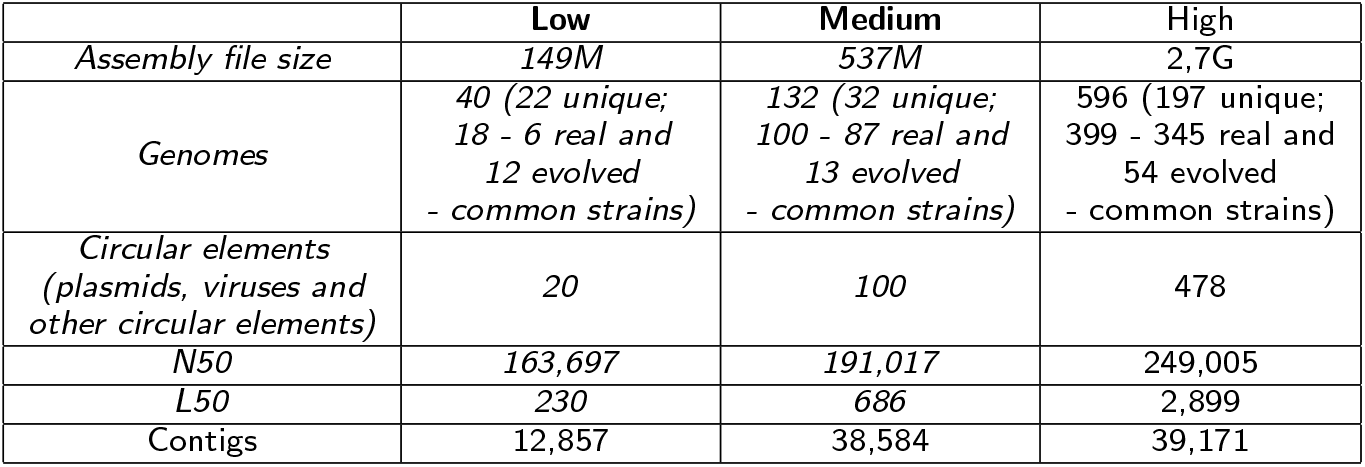
Metagenome assembly statistics from CAMI datasets.

|  | Low | Medium | High |
| --- | --- | --- | --- |
| Assembly file size | 149M | 537M | 2,7G |
| Genomes | 40 (22 unique;<br>18 - 6 real and<br>12 evolved<br>- common strains) | 132 (32 unique;<br>100 - 87 real and<br>13 evolved<br>- common strains) | 596 (197 unique;<br>399 - 345 real and<br>54 evolved<br>- common strains) |
| Circular elements<br>(plasmids, viruses and<br>other circular elements) | 20 | 100 | 478 |
| N50 | 163,697 | 191,017 | 249,005 |
| L50 | 230 | 686 | 2,899 |
| Contigs | 12,857 | 38,584 | 39,171 |

From each assembly, all ORFs were extracted and submitted to CD-HIT [21], a tool for clustering similar sequences, returning the most significant sequences. The low and medium complexity sequences returned, approximately 600,000 sequences, were submitted to InterproScan [22], a tool that searches for protein signatures in different databases (Gene3D, PANTHER, Pfam, PIRSF, PRINTS, ProDom, PROSITE, SMART, SUPERFAMILY and TIGRFAMs). In the high complex sequences, more than 2 billion sequences were returned by CD-HIT. Because of the very high computational costs to classify those sequences using InterproScan, 3 random samples without repetition having 200,000 sequences each were selected. These samples of sequences of high complexity were submitted to InterproScan using the same methodology previously described.

After the identification of sequences by InterproScan, sequences that had annotations found in at least one bank and had IPR annotations (InterPro identifier) were classified as genes and the remaining ones as intergenic. The sequences, InterproScan annotations and their respective classifications (gene or intergenic region) can be found in the supplementary material. The number of positive examples (proteins found by InterproScan), negative examples, and total sequences for each complexity are shown in Table 2. All these datasets are freely available as a new benchmark, being, as far as we are concerned, the largest one available that presents solid ground truth of potential metagenomic genes. For information about the sequence distribution of datasets, see the supplementary material.

**Table 2.** Benchmark dataset using CAMI genome assemblies.

|  | Positive | Negative | Total |
| --- | --- | --- | --- |
| **Low** | 41,068 | 214,521 | 255,589 |
| **Medium** | 57,894 | 289,748 | 347,642 |
| **High (sample 01)** | 34,640 | 165,360 | 200,000 |
| **High (sample 02)** | 34,445 | 165,555 | 200,000 |
| **High (sample 03)** | 34,486 | 165,514 | 200,000 |

For test datasets, the genomes names, the taxonomy ID and the taxonomy level can be found in Table 4 to 9 of the supplementary material.

### Feature Engineering

Several genomic information has been used to build gene predictors, including GC content, sequence length, and others. The GC content corresponds to the percentage of guanine and cytosine bases present in a sequence, being traditionally used in applications to classify genes, since in some cases the coding sequences have higher GC content than non-coding sequences [23]. The length of the sequence is a feature that has the ability to distinguish coding sequences from non-coding ones — it is important in this context because sequences from intergenic regions are, usually, smaller in comparison to the ones found in coding regions [24]. The K-mer frequencies correspond to the number of occurrences of each k-length fragment of a DNA sequence [25]. Codon usage bias refers to the differences in the number of synonymous codons in coding DNA. A codon is a nucleotide triplet that encodes an amino acid (e.g. ATG). Since 64 combinations can be made with 4 nucleotides taken three at a time and considering that there are only 20 amino acids, there is more than one codon per amino acid, in most cases. Two or more codons that encode the same amino acid are called synonymous codon [26], [27].

In previous work, we select 15 features to build the first version of geneRFinder [28]. After feature redundancy evaluation, 11 features were experimentally selected based on the importance index of each feature to the model and the correlations among them. Of these, 4 correspond to GC content, (a) GC content throughout the sequence, (b) GC content from the first position, (c) GC content from the second position, and (d) GC content from the third position of each nucleotide triplet. Another 6 features corresponding to the k-mer frequency, being the frequency variances from 2-mer to 6-mer and the codon usage bias of each of the synonymous codon (c_weight) [29]. Lastly, the sequence length was considered into the feature set. The features have a strong correlation, grouping into two main sets, as shown in Figure 2. The first group refers to GC content features, these features are classic ones in gene prediction, being used in tools such as FragGeneScan, Prodigal and Orphelia. The second group refers to k-mer features, these features are widely used in other branches of Bioinformatics such as assembly [30] and binning [31], but still little explored in gene prediction problems.

**Figure 2.**
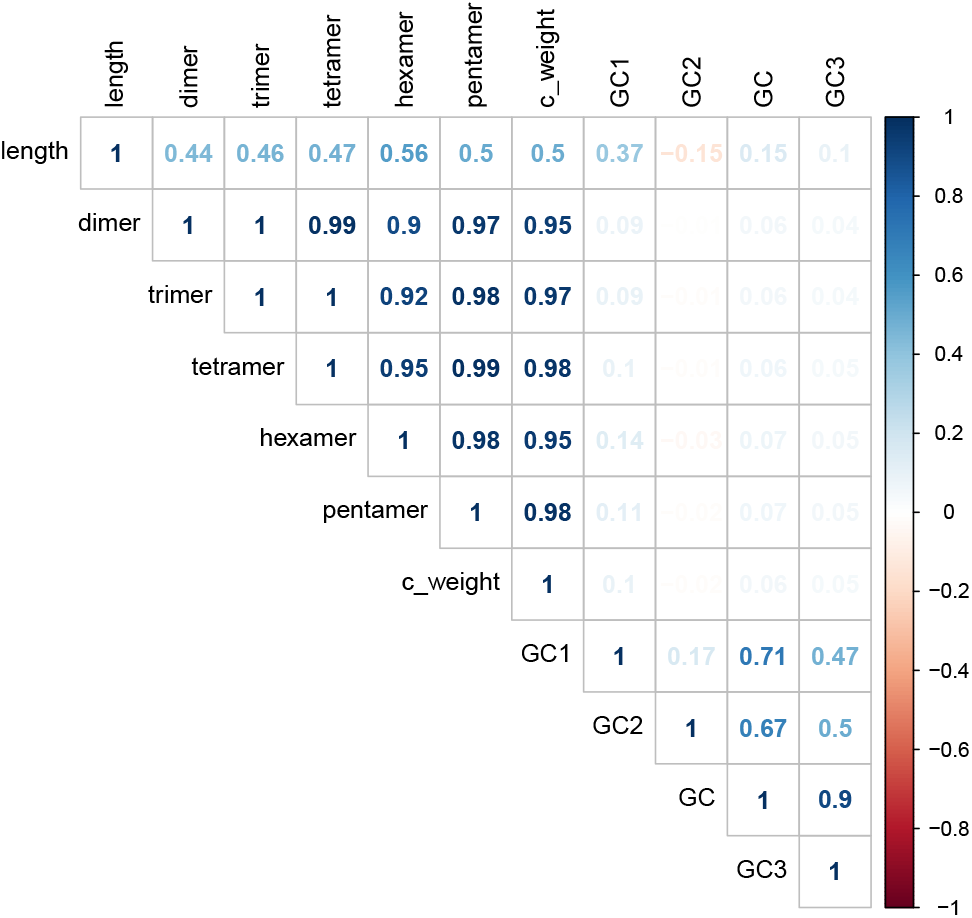
Feature correlation map. Interestingly, k-mer features do not present strong correlation to classical GC content ones.

The feature importance index was calculated according to importance method of the Caret package [32] and, as Figure 3 presents the sequence length as the most important one, followed by k-mer features, having more than 80% importance index. Although GC content features are widely used to discriminate between gene and intergenic regions, in our model they were of minor importance when compared to other features. However, their use in combination with other features influenced the prediction performance, as noted in our previous work [28].

**Figure 3.**
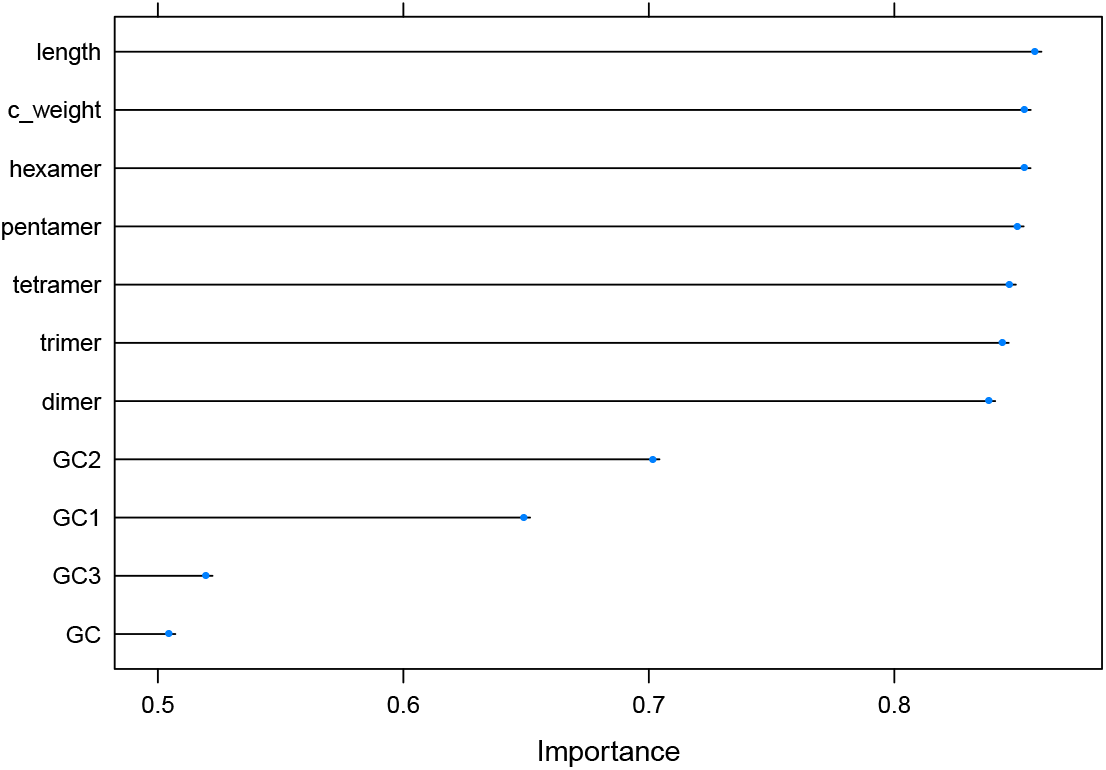
Feature importance plot. K-mer features are more informative than GC content ones.

### Random Forest parametrization

The Random Forest (RF) classifier [33] was used to build the gene prediction model, obtaining better performance when compared to other state-of-the-art predictors. The RF method was chosen based on our previous studies [14] and it was used in similar cases with good performance [34] [35]. Four models having 100, 200, 500 and 700 decision trees with 5-fold cross-validation with 5 repetitions on the performance evaluation training set were built (Figure 4). As stated previously, each instance of the model, which corresponds to a sequence, is represented by 11 numeric features and their respective class. The 700 decision trees model present the best performance result, reaching 92.75% hits on mtry of 3 (number of features considered in each tree node during its construction) for training the geneRFinder model. However, the 100 decision trees model got a similar performance, having less complexity [36], it was the model selected for the geneRFinder tool.

**Figure 4.**
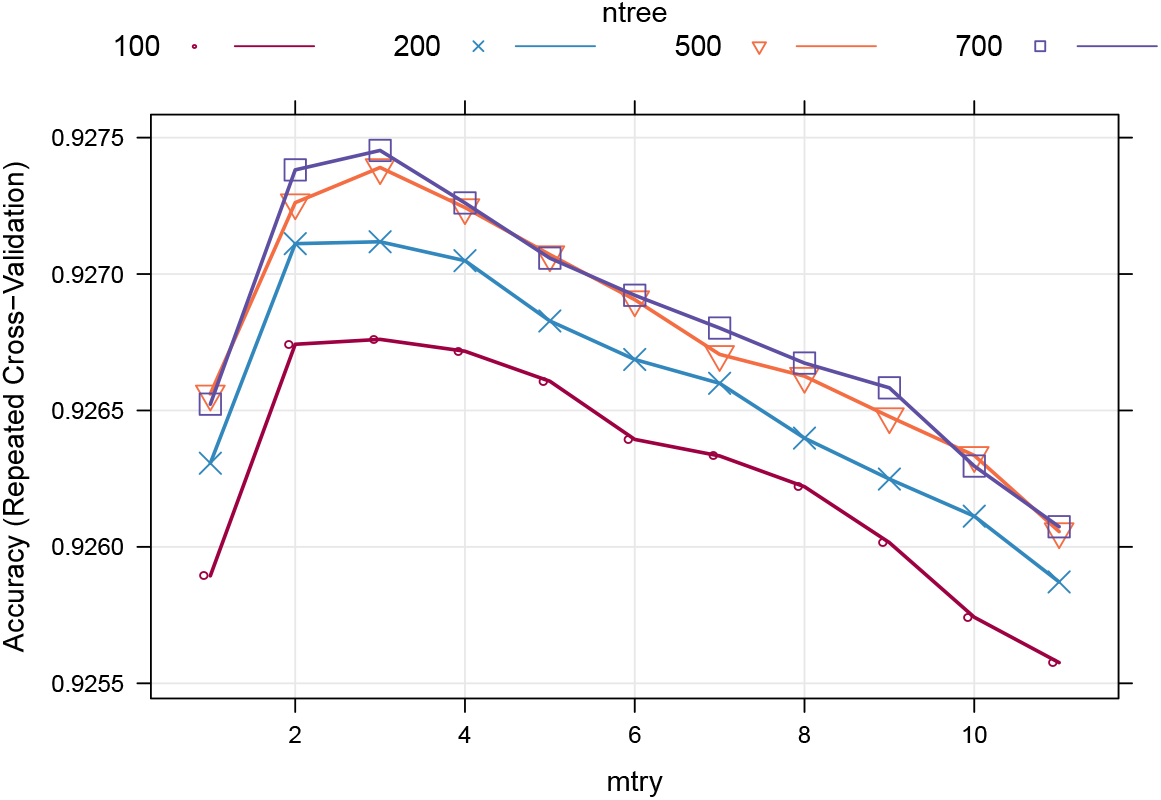
Random Forest trees with 100 models are less complex and robust than large ensembles trees of 700 models.

### Model performance metrics

To evaluate the performance of the gene predictors, four metrics were adopted: accuracy, sensitivity, specificity and AUC. All these metrics express the relations between True/False Positives and True/False Negatives. True Positives are positive examples that were correctly predicted as positive; True Negatives are negative examples that were predicted as negative; False Positives are negative examples that were wrongly predicted as positive; and False Negatives are positive examples that were predicted as negative.

Accuracy represents the hit rate considering the total number of dataset instances and can be defined by the Equation 1.

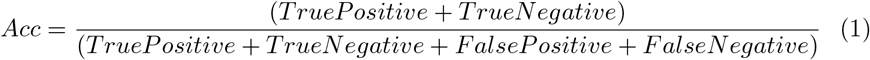

The sensitivity expresses the proportion of annotated genes that have been correctly predicted, and, on the other hand, specificity indicates the percentage of correctly classified intergenic sequences. These measurements are given, respectively, by Equations 2 and 3.

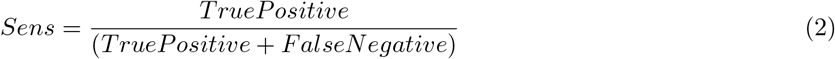

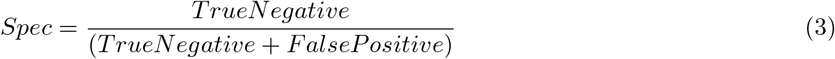

Additionally, AUC (Area Under ROC Curve) [37] is a summary metric that incorporates specificity and sensitivity into a single value.

### The libraries, inputs, outputs and running time

The geneRFinder predictor was built using the R version 3.4.4 [38] language, using the SeqinR package version 3.6-1 [29] for reading sequences and extracting GC content features and the K-mer package version 1.1.2 [39] for extracting k-mer variance features. Model training and sequence prediction are performed using the Caret package version 6.0-84 [32]. The code is parallelized using the doParallel package version 1.0.15 [40]. The predictor allows the user to define triplets considered as start codon (ATG / GTG / TTG), however, in all cases TAA / TAG / TGA will always be considered as a stop codon. The user can also define how many cores can be used by the program. It must be informed as an input parameter to the predictor of the FASTA file containing the reads or contigs to be analyzed. As output, a FASTA file containing the CDS found in the input file is produced, a FASTA file containing the intergenic sequences is optional.

geneRFinder makes predictions at approximately 500 kb/min, or 1000 sequences with 500bp per minute, using 4GB of memory and 5 cores. All scripts and datasets used in this manuscript can be found at https://osf.io/g4qk5/ to reproduce the tests. The geneRFinder tool is freely available at https://github.com/railorena/geneRFinder.

## Results

### Benchmark data

The impact of metagenomic sample complexity on gene prediction was not fully explored by prediction tools until now. There is still no consensus on the datasets used to exploit fair performance comparison of gene prediction tools. Thus, each tool considers different datasets for its analysis.

Although these previous predictors produced similar results, the databases used to evaluate two well-known gene prediction tools - FragGeneScan and Prodigal, for example, contain less than 25% of common organisms (Figure 1). The utilization of specific databases per gene predictor may be justified by the lack of a consolidated benchmark dataset for this purpose.

As with many computational methods, the use of different inputs for gene prediction tools - that is, prediction performance testing using specific organisms - can directly impact the quality of the results produced by these tools, favoring some of them and impact negatively in others. Sequence hit rates can vary considerably for different organisms, due to the sequence similarities found with the training datasets used. Therefore, the performance rates obtained by different tools using different datasets may be biased, making the comparison process between them questionable.

It is evident that the establishment of a standard dataset for metagenomic gene prediction becomes fundamental for improving evaluation of gene predictors. The CAMI challenge paves the direction to make fair benchmarking available to the community but gene prediction was not tackled at that time. In this context, the CAMI-oriented datasets built and used in this work were compiled to provide the scientific community with a fair gene prediction benchmark, ready to use and freely available at https://sourceforge.net/p/generfinder-benchmark.

The benchmark is made up of 9 datasets, as shown in Table 3. For each one of them is provided:

- List of names, taxonomy ID and taxonomic level of the genomes of the organisms that make up the dataset (genomes.csv)
- Set of sequences extracted from the respective selected genomes (sequences.fasta)
- Ground truth for each of the extracted sequences (groundtruth.csv)

**Table 3.** Benchmark description.

| Dataset name | genomes | sequences | CDS | Description |
| --- | --- | --- | --- | --- |
| training1 | 20 | 108,004 | 54,002 | First training set |
| validation | 5 | 19,337 | 14,016 | Validation set used to setup parameters of model built |
| training2 | 129 | 712,886 | 356,443 | Second training set used to build final model |
| test1 | 12 | 54,980 | 31,507 | First test set used to evaluate geneRFinder |
| test2low | 40 | 255,589 | 41,068 | Data extracted from low complexity metagenomic (CAMI) |
| test2medium | 132 | 347,642 | 57,894 | Data extracted from medium complexity metagenomic (CAMI) |
| test2high1 | 160** | 200,000 | 34,640 | Data extracted from high complexity metagenomic (CAMI) (sample 01) |
| test2high2 | 156** | 200,000 | 34,445 | Data extracted from high complexity metagenomic (CAMI) (sample 02) |
| test2high3 | 157** | 200,000 | 34,486 | Data extracted from high complexity metagenomic (CAMI) (sample 03) |

\* the estimated number of genomes was obtained by taxonomic analysis performed by the Kaiju tool [41]

The data provided in the benchmark can be downloaded directly from the browser or using the multiplatform client interface, also available from the benchmark website, through the command line given below, where dataset_name is the database name (training1, training2, etc.) and resource corresponds to the desired file and can assume the values genomes, sequences, ground truth and all - in the last case, to download all the contents of the database.

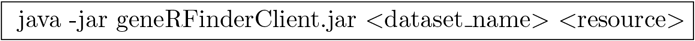

### Revisiting gene prediction

To analyze the geneRFinder performance, prediction tests were performed based on sequence length, being a fundamental feature, the most important feature in our model, to discriminate whether an sequence is coding or not. We performed the tests in dataset test1 (with 12 genomes) to predict sequences from 100bp to 2000bp, as presented in Figure 5. geneRFinder achieved accuracy and sensitivity performance above 75% for sequences above 200bp, reaching more than 90% for sequences above 500bp. The specificity of geneRFinder reached more than 75% for all sequences, reaching a higher percentage in larger sequences. This test showed that the longer the sequence, the more information there is to characterize it, but even in small sequences, its performance was satisfactory.

**Figure 5.**
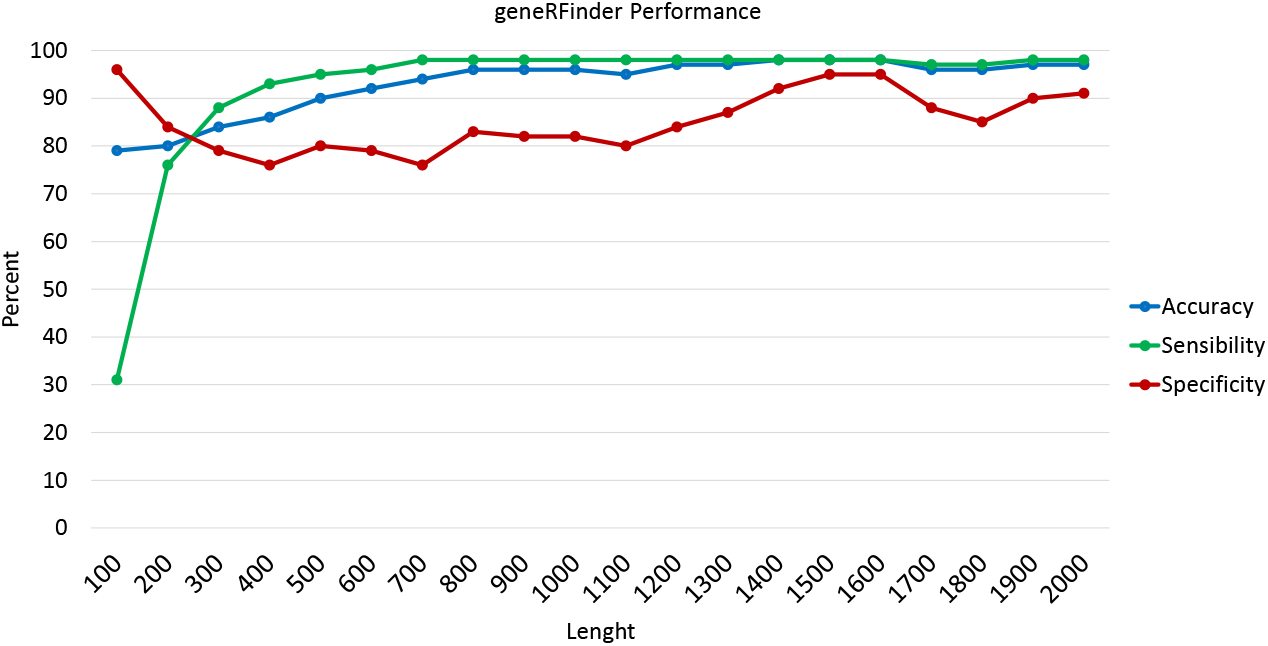
geneRFinder performance in sequences of different lengths using the dataset test1.

We used FragGeneScan and Prodigal for comparison analysis using the introduced benchmark. FragGeneScan is considered one of the best performing tools for gene prediction [42], being used by EBI Metagenomics [43] and MG-RAST [44], two important pipelines for metagenomic data analysis [45]. Prodigal was added to the EBI Metagenomics pipeline as a complement to FragGeneScan to predict large sequences [43], while only FragGeneScan is used for small sequences.

For the dataset test1 (with 12 genomes), the prediction results are shown in Figure 6. The best accuracy was obtained with the geneRFinder, with a percentage difference of approximately 20% more than FragGeneScan and Prodigal. With less considerable distances than the others, the best sensitivity was obtained with FragGeneScan, with differences of 2% and 7% against Prodigal and geneRFinder, respectively. The geneRFinder also achieved better specificity performance, with 55 percentage points higher than Prodigal and 60 percentage points higher than FragGeneScan.

**Figure 6.**
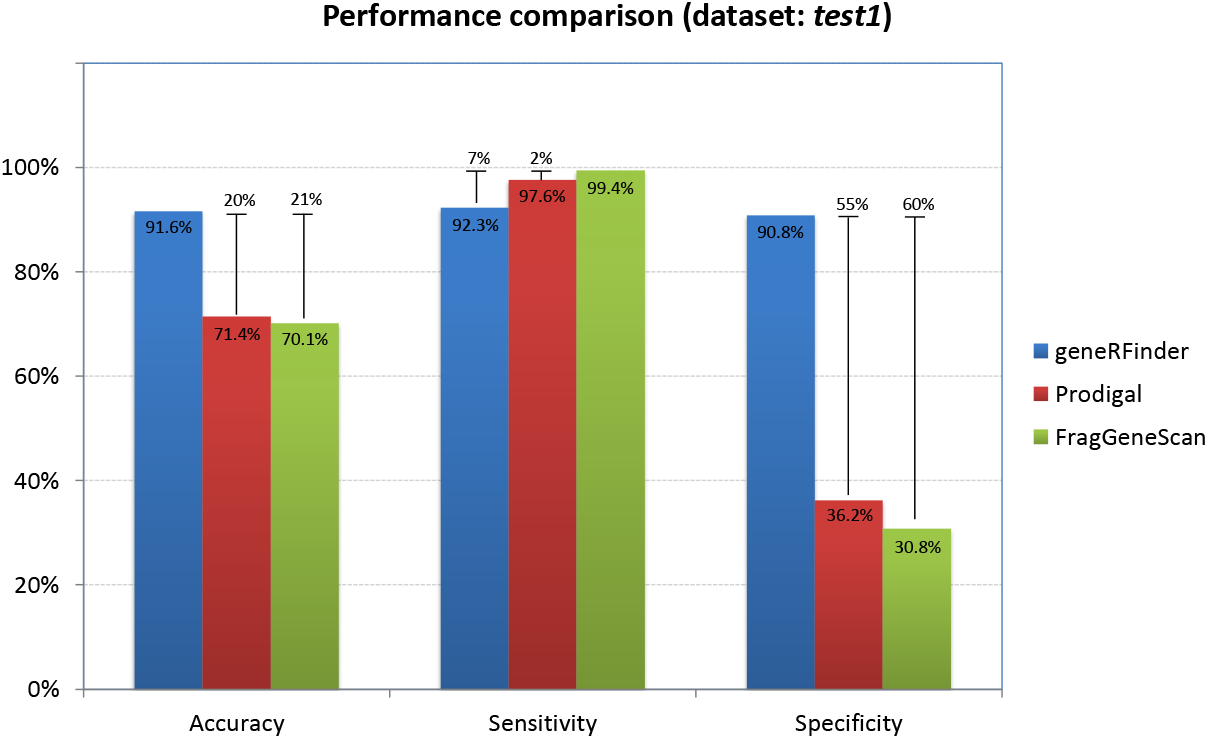
Gene predictors performance using the dataset test1.

When evaluating predictors performance in sequences from low complexity metagenome (test2low), Figure 7, geneRFinder obtained the best accuracy, with a percentage variation of 53% compared to Prodigal and 63% against FragGeneScan. In sensitivity, FragGeneScan hit 99% of the data, 1% more than Prodigal and 5% more than geneRFinder. In geneRFinder specificity obtained better result, correctly classifying 64% more sequences than Prodigal and 76% more than FragGeneScan.

**Figure 7.**
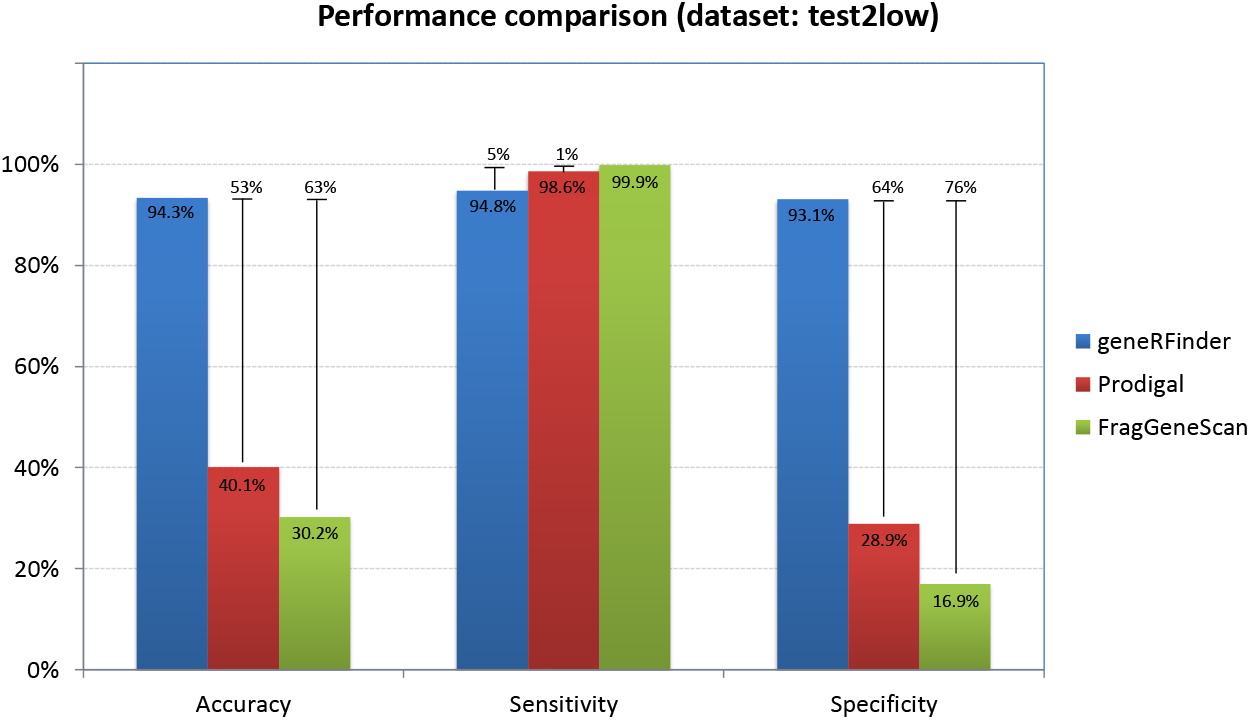
Gene predictors performance using the low complexity metagenome.

In the results of medium complexity sequences (test2medium), Figure 8, the accuracy of the RFinder gene was 93.8%, compared to 40.1% for Prodigal and 29.6% for FragGeneScan. FragGeneScan sensitivity differed in percentage terms by only 2% of Prodigal and 5% of geneRFinder. In specificity geneRFinder again had better performance (93.6%), with 65 percentage points more than Prodigal and 78 more than FragGeneScan.

For the tests in the high complexity metagenome, the predictions were made in the 3 datasets (test2high1, test2high2 and test2high3) and the average of the results is presented in Figure 9. geneRFinder achieved better performance in accuracy, with 93.4% correctness and percentage differences of 54% and 64%, respectively, against Prodigal and FragGeneScan. In sensitivity, FragGeneScan showed 2 percentage points higher than Prodigal and 5 percentage points higher than geneRFinder. In specificity, geneRFinder had the largest distance against FragGeneScan, 79 percentage points, and 66 more than Prodigal.

**Figure 8.**
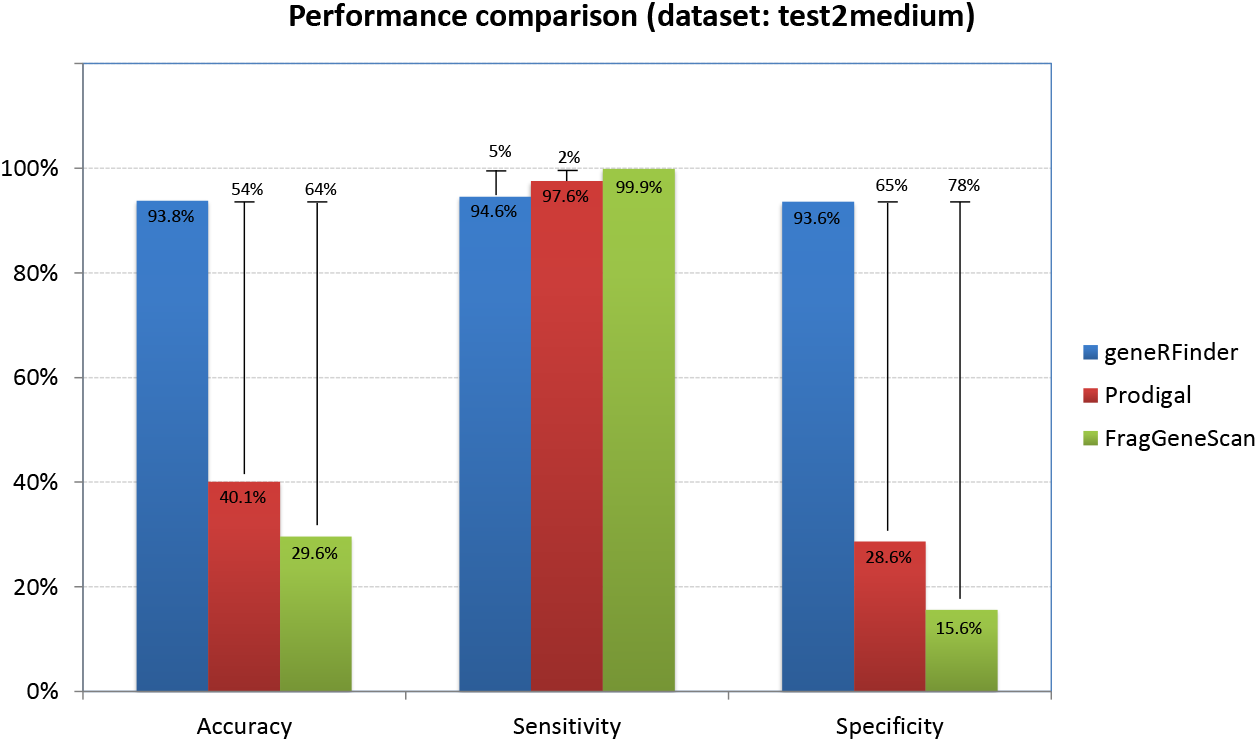
Gene predictors performance using the medium complexity metagenome.

**Figure 9.**
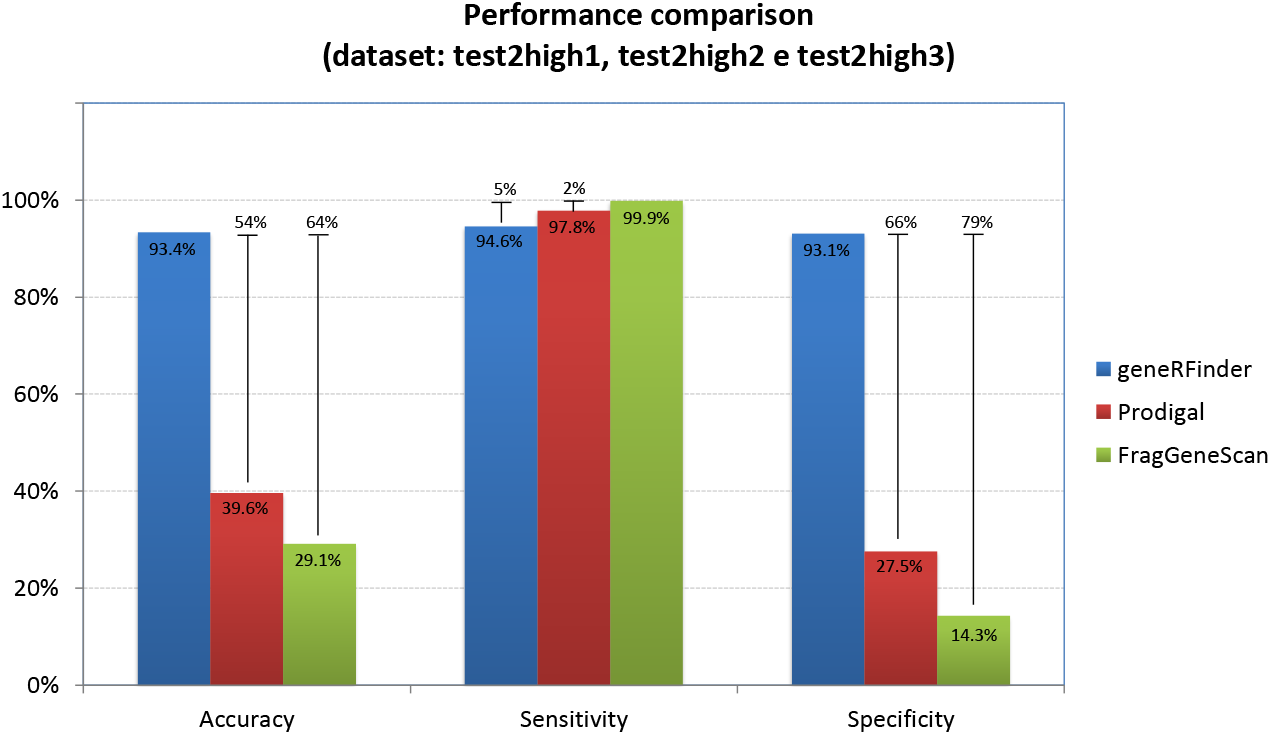
Gene predictors performance using the high complexity metagenomes.

FragGeneScan and Prodigal had difficulty to discriminate which sequences were not coding ones, on the other hand, geneRFinder was able to more clearly detect both coding and non-coding sequences, reaching more than 90% specificity in all datasets. The high specificity of geneRFinder allows predicting without requiring other segments of the sequence, such as translation initiation site, since its features capture the signals present in coding and non-coding sequences independently.

When analyzing the proportion of predictors sensitivity and specificity represented by the ROC curve in Figure 10, geneRFinder achieved better performance in the 4 datasets. This proportion, measured as a percentage by the area under the ROC curve, was at least 24 percentage points higher than in other tools. The individual predictions of each dataset can be found in the supplementary material.

**Figure 10.**
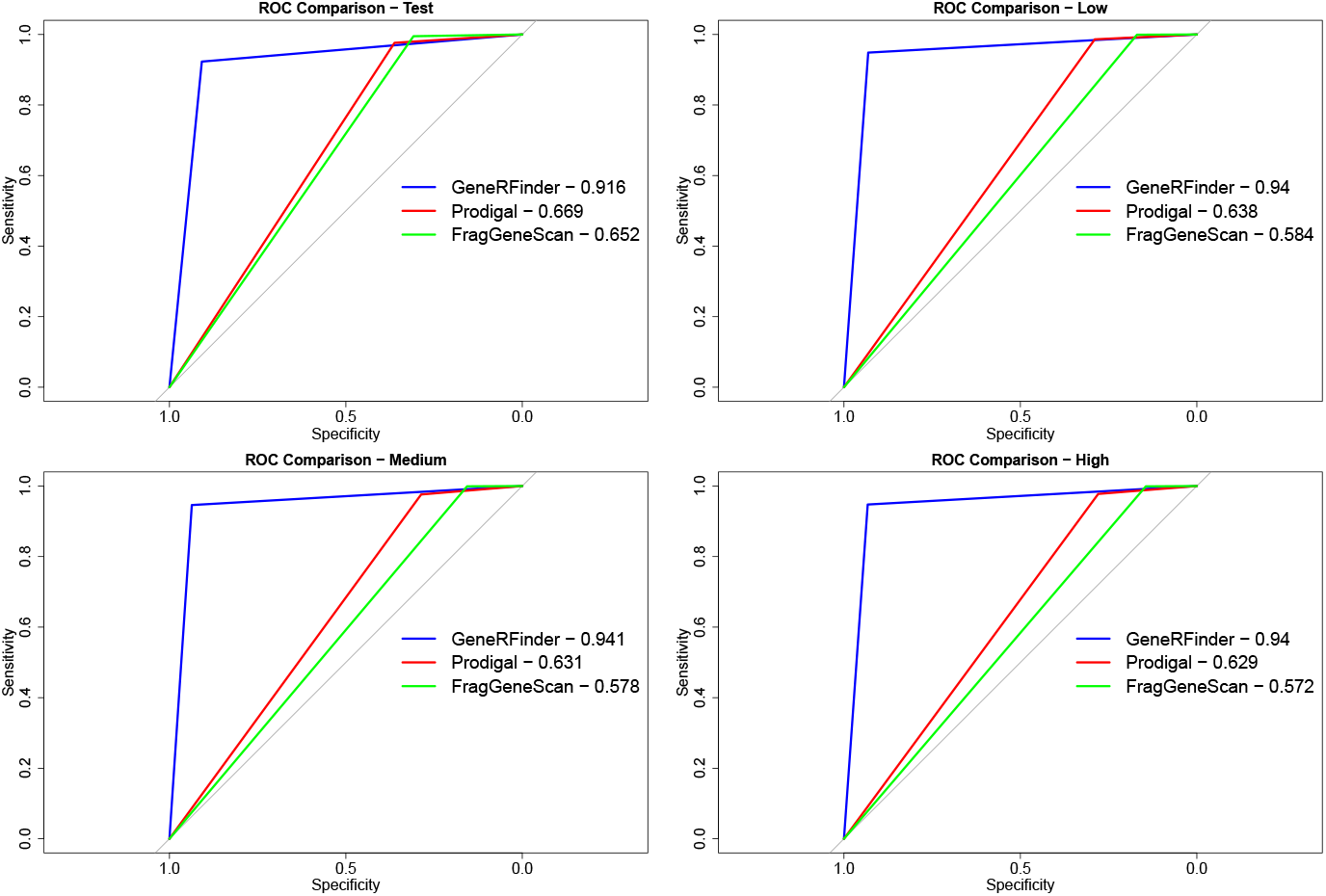
ROC curves for gene predictors using the benchmark.

In order to verify if the differences in prediction performances of the geneRFinder, FragGeneScan, and Prodigal were significant, we calculated the statistical differences between predictors performance using McNemar’s test [46] for all datasets. McNemar’s test is suitable for comparing the performances of distinct ML classifiers, especially when it is not possible or feasible to train models several times, as is the case with FragGeneScan and Prodigal tools, which are available with ML models previously trained.

McNemar’s tests were performed for all possible pair combinations between predictors and performed individually on each complexity. Unanimously, the null hypothesis - that both predictors show equivalent performances - was rejected with a 99% statistical confidence interval, thus indicating that the superiority of geneRFinder over the other tools results from statistically effective differences, and not just casual variations of performance. The codes and input files needed to reproduce these tests can be found at https://osf.io/g4qk5/ and the contingency tables can be found in Table 10 to Table 15 of the supplementary material.

The accuracy, sensitivity and specificity express the performance of the classifier under different perspectives; as mentioned previously, the accuracy presents the general success rate of the classifier, but it does not offer the conditions to evaluate the strengths and weaknesses of each one of them; the sensitivity and specificity, respectively, allow us to analyze the performance of the classifier by predicting sequences known to be positive (corresponding to CDS) and negative. FragGeneScan, for example, has the best average sensitivity rate. This means that, from all sequences that corresponded to CDS, this tool rated approximately 99% of them correctly on all datasets. However, this same tool misclassified an average of 80% of non-CDS sequences. In gene annotation processes, in which experts perform the painstaking work of trying to identify the gene corresponding to each CDS, according to these statistics, many sequences will undergo such annotation. Low specificity in this context implies in undue submission of several sequences to the annotation process and, consequently, in the waste of working time. In contrast, geneRFinder could achieve superior rates for specificity — beyond equivalent rates for sensitivity, what can be seen in Figure 9, demonstrating its general superiority.

## Conclusion

Gene prediction is a classical and key challenge in (meta)genomics. Computational methods for gene finding are mostly based on machine learning strategies. In the very beginning gene predictor explored the power of Hidden Markov Models, evolving to the exploration of neural networks, support vectors and recently ensemble strategies (Random Forest, Gradient Boosting Machines, etc.). Gene predictors usually provided similar results though they differ clearly in their benchmark data. Thus, there is some skepticism regarding the extent to which the model’s performance of these gene predictors was fairly taken into account during comparison analysis. This situation is more critical in large scale and complex biological datasets like those in metagenomics.

We provided a new benchmark data based on the well-known CAMI challenge. CAMI provides datasets of unprecedented complexity and degree of realism, though it does not provide datasets to assess gene predictors. We generate nine datasets of distinct complexities, being 5 of them derived from available CAMI metagenome assemblies to assess the robustness of gene predictors, making it freely available for future benchmarking, and the remaining 4 datasets manually developed.

The geneRFinder is introduced to deal with the prediction of protein-coding in distinct metagenomic complexities. Comparison analysis with state-of-the-art gene predictors highlights its utility, providing a good balance between sensitivity and specificity performance metrics.

