## supplementary material for "geneRFinder: gene finding in distinct metagenomic data complexities"

1. Additional information about GeneRFinder-Benchmark

The ORFs extracted from CAMI genome assemblies have similar size distribution, with a predominance of sequences smaller than 200bp (Figure1). This is due to the fact that ORF extraction considered as ORF all sequences started with start codon and ended with some stop codon, regardless of its position. In this way, ORFs that were inside ORFs were also extracted. Thus, even though an ORF is part of the coding sequence, does not represent the entire gene, but only a fraction of it.

ORFs from the 12 annotated genomes of the first set of tests, named Test in Figure1, have a more balanced size distribution, however, ORFs under 200bp are also greater in this dataset. It is worth noting that the amount of ORFs less than 200bp is inversely proportional to the complexity of these samples. This is probably due to the fact that low complexity samples have, by definition, fewer genomes [1], that can reduce the complexity of the assembly process and thus obtaining larger and more reliable sequences, which will be used for the process of ORF extraction. This scenario may justify the increase in ORFs as complexity decreases.

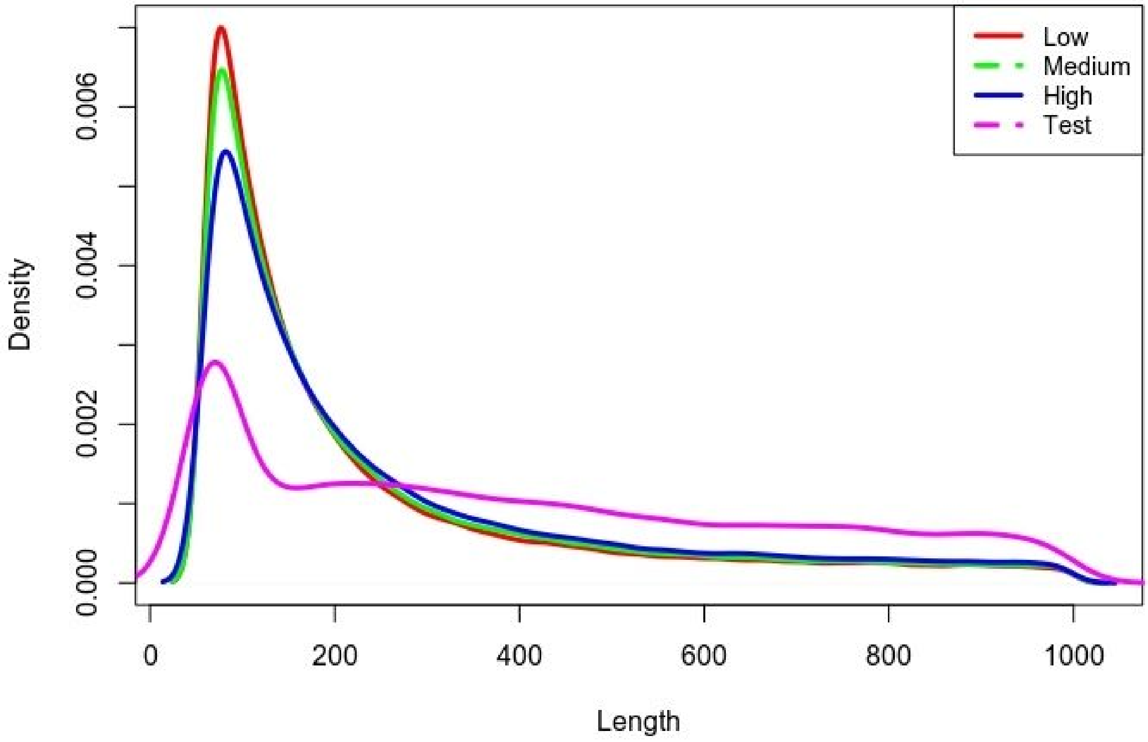


Figure 1. ORFs length distribution in the benchmark dataset.

2. Additional information about simulated genomes from CAMI

In order to increase strain level diversity, CAMI used 12 strain genomes that were simulated to produce the low complexity dataset. All information about related organisms and simulation process can be found at CAMI paper [2]. The names of the organisms of the simulated genomes mentioned are presented in the supplementary tables 5 to 9, using the same nomenclature of the genome files provided by CAMI.

Supplementary Table 1. List of genomes used in the initial training (training1)

| Taxonomy_ID | Name | Rank |
|---|---|---|
| 62977 | Acinetobacter sp ADP1 | species |
| 272557 | Aeropyrum pernix | species |
| 264462 | Bdellovibrio bacteriovorus | species |
| 257310 | Bordetella bronchiseptica | species |
| 262698 | Brucella abortus 9-941 | species |
| 195099 | Campylobacter jejuni RM1221 | species |
| 203907 | Candidatus Blochmannia floridanus | species |
| 159087 | Dechloromonas aromatica RCB | species |
| 243164 | Dehalococcoides ethenogenes 195 | species |
| 226185 | Enterococcus faecalis V583 | species |
| 177416 | Francisella tularensis tularensis | species |
| 106370 | Frankia casuarinae | species |
| 251221 | Gloeobacter violaceus | species |
| 290633 | Gluconobacter oxydans 621H | species |
| 233412 | Haemophilus ducreyi 35000HP | species |
| 349521 | Hahella chejuensis KCTC 2396 | species |
| 283942 | Idiomarina loihiensis L2TR | species |
| 272621 | Lactobacillus acidophilus NCFM | species |
| 272623 | Lactococcus lactis | species |
| 265311 | Mesoplasma florum L1 | species |

Supplementary Table 2. List of genomes used in the validation (validation)

| Taxonomy_ID | Name | Rank |
|---|---|---|
| 348780 | Natronomonas pharaonis | species |
| 107806 | Buchnera aphidicola | species |

| | | |
|---|---|---|
| 272560 | Burkholderia pseudomallei | species |
| 224308 | Bacillus subtilis | species |
| 306537 | Corynebacterium jeikeium | species |

Supplementary Table 3. List of genomes used in the full training (training2)

| Taxonomy_ID | Name | Rank |
|---|---|---|
| 272569 | Haloarcula marismortui ATCC 43049 chromosome I | species |
| 272569 | Haloarcula marismortui ATCC 43049 chromosome II | species |
| 64091 | Halobacterium salinarum NRC-1 | species |
| 267377 | Methanococcus maripaludis S2 | species |
| 188937 | Methanosarcina acetivorans C2A | species |
| 339860 | Methanosphaera stadtmanae DSM 3091 | species |
| 187420 | Methanothermobacter thermautotrophicus str. Delta H | species |
| 263820 | Picrophilus torridus DSM 9790 | species |
| 330779 | Sulfolobus acidocaldarius DSM 639 | species |
| 69014 | Thermococcus kodakarensis KOD1 | species |
| 273075 | Thermoplasma acidophilum DSM 1728 | species |
| 62977 | Acinetobacter sp. ADP1 | species |
| 176299 | Agrobacterium tumefaciens str. C58 chromosome circular | species |
| 176299 | Agrobacterium tumefaciens str. C58 chromosome linear | species |
| 234826 | Anaplasma marginale str. St. Maries | species |
| 224324 | Aquifex aeolicus VF5 | species |
| 322098 | Aster yellows witches'-broom phytoplasma AYWB | species |
| 76114 | Azoarcus sp. EbN1 | species |
| 272559 | Bacteroides fragilis NCTC 9343 | species |
| 283166 | Bartonella henselae str. Houston-1 | species |
| 264462 | Bdellovibrio bacteriovorus HD100 | species |

| | | |
|---|---|---|
| 206672 | Bifidobacterium longum NCC2705 | species |
| 257310 | Bordetella bronchiseptica RB50 | species |
| 224326 | Borrelia burgdorferi B31 | species |
| 224911 | Bradyrhizobium japonicum USDA 110 | species |
| 262698 | Brucella abortus biovar 1 str. 9-941 chromosome I | species |
| 262698 | Brucella abortus biovar 1 str. 9-941 chromosome II | species |
| 195099 | Campylobacter jejuni RM1221 | species |
| 203907 | Candidatus Blochmannia floridanus | species |
| 246194 | Carboxydothermus hydrogenoformans Z-2901 | species |
| 190650 | Caulobacter crescentus CB15 | species |
| 243161 | Chlamydia muridarum Nigg | species |
| 218497 | Chlamydophila abortus S26/3 | species |
| 243365 | Chromobacterium violaceum ATCC 12472 | species |
| 272562 | Clostridium acetobutylicum ATCC 824 | species |
| 167879 | Colwellia psychrerythraea 34H | species |
| 227377 | Coxiella burnetii RSA 493 | species |
| 321327 | Cyanobacteria bacterium Yellowstone A-Prime | species |
| 159087 | Dechloromonas aromatica RCB | species |
| 243164 | Dehalococcoides ethenogenes 195 | species |
| 243230 | Deinococcus radiodurans R1 | species |
| 177439 | Desulfotalea psychrophila LSv54 | species |
| 207559 | Desulfovibrio desulfuricans G20 | species |
| 269484 | Ehrlichia canis str. Jake | species |
| 226185 | Enterococcus faecalis V583 | species |
| 218491 | Erwinia carotovora subsp. atroseptica SCRI1043 | species |
| 177416 | Francisella tularensis subsp. tularensis SCHU S4 | species |
| 106370 | Frankia casuarinae | species |

| | | |
|---|---|---|
| 190304 | Fusobacterium nucleatum subsp. nucleatum ATCC 25586 | species |
| 235909 | Geobacillus kaustophilus HTA426 | species |
| 269799 | Geobacter metallireducens GS-15 | species |
| 251221 | Gloeobacter violaceus PCC 7421 | species |
| 290633 | Gluconobacter oxydans 621H | species |
| 233412 | Haemophilus ducreyi 35000HP | species |
| 349521 | Hahella chejuensis KCTC 2396 | species |
| 283942 | Idiomarina loihiensis L2TR | species |
| 272621 | Lactobacillus acidophilus NCFM | species |
| 272623 | Lactococcus lactis subsp. lactis Il1403 | species |
| 297245 | Legionella pneumophila str. Lens | species |
| 281090 | Leifsonia xyli subsp. xyli str. CTCB07 | species |
| 267671 | Leptospira interrogans serovar Copenhageni str. Fiocruz L1-130 chromosome I | species |
| 267671 | Leptospira interrogans serovar Copenhageni str. Fiocruz L1-130 chromosome II | species |
| 272626 | Listeria innocua Clip11262 | species |
| 342108 | Magnetospirillum magneticum AMB-1 | species |
| 221988 | Mannheimia succiniciproducens MBEL55E | species |
| 265311 | Mesoplasma florum L1 | species |
| 266835 | Mesorhizobium loti MAFF303099 | species |
| 243233 | Methylococcus capsulatus str. Bath | species |
| 264732 | Moorella thermoacetica ATCC 39073 | species |
| 262316 | Mycobacterium avium subsp. paratuberculosis K-10 | species |
| 340047 | Mycoplasma capricolum subsp. capricolum ATCC 27343 | species |
| 242231 | Neisseria gonorrhoeae FA 1090 | species |
| 323098 | Nitrobacter winogradskyi Nb-255 | species |
| 323261 | Nitrosococcus oceani ATCC 19707 | species |

| | | |
|---|---|---|
| 228410 | Nitrosomonas europaea ATCC 19718 | species |
| 323848 | Nitrosospira multiformis ATCC 25196 | species |
| 247156 | Nocardia farcinica IFM 10152 | species |
| 279238 | Novosphingobium aromaticivorans DSM 12444 | species |
| 221109 | Oceanobacillus iheyensis HTE831 | species |
| 262768 | Onion yellows phytoplasma OY-M | species |
| 272843 | Pasteurella multocida subsp. multocida str. Pm70 | species |
| 338963 | Pelobacter carbinolicus DSM 2380 | species |
| 319225 | Pelodictyon luteolum DSM 273 | species |
| 298386 | Photobacterium profundum SS9 | species |
| 298386 | Photobacterium profundum SS9 chromosome 2 | species |
| 243265 | Photorhabdus luminescens subsp. laumondii TTO1 | species |
| 242619 | Porphyromonas gingivalis W83 | species |
| 267747 | Propionibacterium acnes KPA171202 | species |
| 326442 | Pseudoalteromonas haloplanktis TAC125 chromosome I | species |
| 326442 | Pseudoalteromonas haloplanktis TAC125 chromosome II | species |
| 208964 | Pseudomonas aeruginosa PAO1 | species |
| 259536 | Psychrobacter arcticus 273-4 | species |
| 264198 | Ralstonia eutropha JMP134 chromosome 1 | species |
| 264198 | Ralstonia eutropha JMP134 chromosome 2 | species |
| 347834 | Rhizobium etli CFN 42 | species |
| 272943 | Rhodobacter sphaeroides 2.4.1 chromosome 1 | species |
| 272943 | Rhodobacter sphaeroides 2.4.1 chromosome 2 | species |
| 243090 | Rhodopirellula baltica SH 1 | species |
| 258594 | Rhodopseudomonas palustris CGA009 | species |
| 269796 | Rhodospirillum rubrum ATCC 11170 | species |
| 272944 | Rickettsia conorii str. Malish 7 | species |

| | | |
|---|---|---|
| 309807 | Salinibacter ruber DSM 13855 | species |
| 321314 | Salmonella enterica subsp. enterica serovar Choleraesuis str. SC-B67 | species |
| 211586 | Shewanella oneidensis MR-1 | species |
| 300268 | Shigella boydii Sb227 | species |
| 246200 | Silicibacter (ncbi = Ruegeria) pomeroyi DSS-3 | species |
| 266834 | Sinorhizobium meliloti 1021 | species |
| 343509 | Sodalis glossinidius str. 'morsitans' | species |
| 273036 | Staphylococcus aureus RF122 | species |
| 208435 | Streptococcus agalactiae 2603V/R | species |
| 227882 | Streptomyces avermitilis MA-4680 | species |
| 292459 | Symbiobacterium thermophilum IAM 14863 | species |
| 1148 | Synechocystis sp. PCC 6803 | species |
| 273068 | Thermoanaerobacter tengcongensis MB4 | species |
| 269800 | Thermobifida fusca YX | species |
| 197221 | Thermosynechococcus elongatus BP-1 | species |
| 243274 | Thermotoga maritima MSB8 | species |
| 262724 | Thermus thermophilus HB27 | species |
| 292415 | Thiobacillus denitrificans ATCC 25259 | species |
| 317025 | Thiomicrospira crunogena XCL-2 | species |
| 243275 | Treponema denticola ATCC 35405 | species |
| 218496 | Tropheryma whipplei TW08/27 | species |
| 273119 | Ureaplasma parvum serovar 3 str. ATCC 700970 | species |
| 243277 | Vibrio cholerae O1 biovar eltor str. N16961 chromosome I | species |
| 243277 | Vibrio cholerae O1 biovar eltor str. N16961 chromosome II | species |
| 36870 | Wigglesworthia glossinidia endosymbiont of Glossina brevipalpis | species |
| 273121 | Wolinella succinogenes DSM 1740 | species |

| | | |
|---|---|---|
| 190486 | Xanthomonas axonopodis pv. citri str. 306 | species |
| 160492 | Xylella fastidiosa 9a5c | species |

Supplementary Table 4. List of genomes used in the test (test1)

| Taxonomy_ID | Name | Rank |
|---|---|---|
| 243232 | Methanocaldococcus jannaschii | species |
| 224325 | Archaeoglobus fulgidus | species |
| 348780 | Natronomonas pharaonis | species |
| 83332 | Mycobacterium tuberculosis | species |
| 272560 | Burkholderia pseudomallei | species |
| 224308 | Bacillus subtilis subsp. subtilis | species |
| 306537 | Corynebacterium jeikeium | species |
| 194439 | Chlorobium tepidum 0.8475 4 | species |
| 511145 | Escherichia coli str. K-12 | species |
| 85963 | Helicobacter pylori | species |
| 74546 | Prochlorococcus marinus | species |
| 292805 | Wolbachia endosymbiont | species |

Supplementary Table 5. List of genomes used in the low complexity dataset (test2low)

| Taxonomy_ID | Name | Rank |
|---|---|---|
| 1123015 | Pseudomonas aeruginosa | species |
| 1121001 | Andreprevotia lacus | species |
| 1121885 | Flavisolibacter ginsengiterrae | species |
| 1122187 | Lysobacter oryzae | species |
| 644383 | Thermosporothrix hazakensis | species |
| 2088 | Anaeroplasma bactoclasticum | species |
| 1121884 | Flavisolibacter ginsengisoli | species |
| 1123243 | Schwartzia succinivorans | species |

| | | |
|---|---|---|
| 1122156 | Lampropedia hyalina | species |
| 1123349 | Tepidibacter formicigenes | species |
| 1121301 | Paramaledivibacter caminithermalis | species |
| 1120989 | Anaerobranca californiensis | species |
| 1120996 | Anaerosporobacter mobilis | species |
| 1121393 | Desulfatibacillum alkenivorans | species |
| 2371 | Xylella fastidiosa | species |
| 328515 | Nonlabens dokdonensis | species |
| 404881 | Defluviimonas denitrificans | species |
| 1004304 | Hydrotalea sandarakina | species |
| 990712 | Albidovulum xiamenense | species |
| 266 | Paracoccus denitrificans | species |
| 266 | Paracoccus denitrificans | species |
| 266 | Paracoccus denitrificans | species |
| 29580 | Janthinobacterium | genus |
| 173053 | Tetrasphaera duodecadis | species |
| 75309 | Rhodanobacter | genus |
| 2037 | Actinomycetales | order |
| 1385 | Bacillales | order |
| 29580 | Janthinobacterium | genus |
| 2088 | evo_1035930.011 | strain |
| 2088 | evo_1035930.029 | strain |
| 2088 | evo_1035930.032 | strain |
| 1004304 | evo_1049056.011 | strain |
| 1004304 | evo_1049056.013 | strain |
| 1004304 | evo_1049056.015 | strain |
| 1004304 | evo_1049056.031 | strain |

| | | |
|---|---|---|
| 1004304 | evo_1049056.039 | strain |
| 1385 | evo_1286_AP.008 | strain |
| 1385 | evo_1286_AP.026 | strain |
| 1385 | evo_1286_AP.033 | strain |
| 1385 | evo_1286_AP.037 | strain |

Supplementary Table 6. List of genomes used in the medium complexity dataset (test2medium)

| Taxonomy_ID | Name | Rank |
|---|---|---|
| 1763 | Mycobacterium | genus |
| 80864 | Comamonadaceae | family |
| 80864 | Comamonadaceae | family |
| 2062 | Streptomycetaceae | family |
| 1707 | Cellulomonas | genus |
| 1121911 | Garciella nitratireducens | species |
| 1121270 | Carboxydocella sporoproducens | species |
| 1123491 | Vibrio cincinnatiensis | species |
| 1121442 | Desulfovibrio bizertensis | species |
| 1123232 | Salinicoccus kunmingensis | species |
| 1123358 | Tetragenococcus halophilus | species |
| 1120981 | Alysiella filiformis | species |
| 1121478 | Diaphorobacter oryzae | species |
| 1121466 | Desulfovibrio vietnamensis | species |
| 1122961 | Picrophilus oshimae | species |
| 1122958 | Phenylobacterium composti | species |
| 1123266 | Sphingomonas aestuarii | species |
| 1121439 | Desulfovibrio alkalitolerans | species |
| 1123492 | Vibrio gazogenes | species |

| | | |
|---|---|---|
| 39498 | [Eubacterium] yurii | species |
| 745369 | Acetoanaerobium noterae | species |
| 35623 | Acholeplasma oculi | species |
| 915 | Nitrosomonas europaea | species |
| 466 | Legionella maceachernii | species |
| 393921 | Porphyromonas crevioricanis | species |
| 1513 | Clostridium tetani | species |
| 28136 | Prevotella oulorum | species |
| 1122184 | Lutispora thermophila | species |
| 1121088 | Bacillus coagulans | species |
| 1121881 | Ferrithrix thermotolerans | species |
| 1121264 | Caloranaerobacter azorensis | species |
| 1121128 | Butyricicoccus pullicaecorum | species |
| 1121316 | Clostridium grantii | species |
| 1121131 | Butyrivibrio fibrisolvens | species |
| 1121132 | Butyrivibrio hungatei | species |
| 1123350 | Tepidibacter thalassicus | species |
| 1123003 | Propionispora hippei | species |
| 1121428 | Desulfotomaculum hydrothermale | species |
| 1121266 | Caminicella sporogenes | species |
| 445973 | Intestinibacter bartlettii | species |
| 299255 | Ferrimonas marina | species |
| 1123357 | Tessaracoccus bendigoensis | species |
| 1123029 | Pseudoxanthobacter soli | species |
| 103732 | Lechevalieria flava | species |
| 68170 | Lechevalieria aerocolonigenes | species |
| 2030 | Kibdelosporangium aridum | species |

| | | |
|---|---|---|
| 639310 | Olleya aquimaris | species |
| 688913 | Ohtaekwangia kribbensis | species |
| 688867 | Ohtaekwangia koreensis | species |
| 1061 | Rhodobacter capsulatus | species |
| 121821 | Roseinatronobacter thiooxidans | species |
| 441209 | Rhodobaca barguzinensis | species |
| 561061 | Sphingobacterium psychroaquaticum | species |
| 570520 | Formosa spongicola | species |
| 1513896 | Sphingobacterium nematocida | species |
| 354 | Azotobacter vinelandii | species |
| 82367 | Paracoccus pantotrophus | species |
| 82367 | Paracoccus pantotrophus | species |
| 2242 | Halobacterium salinarum | species |
| 2242 | Halobacterium salinarum | species |
| 1061 | Rhodobacter capsulatus | species |
| 759851 | Sporosarcina newyorkensis | species |
| 759851 | Sporosarcina newyorkensis | species |
| 382 | Sinorhizobium meliloti | species |
| 382 | Sinorhizobium meliloti | species |
| 1525 | Moorella thermoacetica | species |
| 1525 | Moorella thermoacetica | species |
| 1082 | Phaeospirillum fulvum | species |
| 1082 | Phaeospirillum fulvum | species |
| 1082 | Phaeospirillum fulvum | species |
| 1082 | Phaeospirillum fulvum | species |
| 1516 | Thermoanaerobacter thermohydrosulfuricus | species |
| 1516 | Thermoanaerobacter thermohydrosulfuricus | species |

| | | |
|---|---|---|
| 192 | Azospirillum brasilense | species |
| 629679 | Promicromonospora umidemergens | species |
| 280 | Xanthobacter autotrophicus | species |
| 69278 | Aquamicrobium | genus |
| 85413 | Bosea | genus |
| 379 | Rhizobium | genus |
| 13687 | Sphingomonas | genus |
| 80864 | Comamonadaceae | family |
| 162491 | Oerskovia | genus |
| 80864 | Comamonadaceae | family |
| 1707 | Cellulomonas | genus |
| 80864 | Comamonadaceae | family |
| 13687 | Sphingomonas | genus |
| 169973 | uncultured Massilia sp. | species |
| 1143711 | Paenibacillus frigoriresistens | species |
| 367298 | Phycicoccus | genus |
| 1707 | Cellulomonas | genus |
| 85023 | Microbacteriaceae | family |
| 13687 | Sphingomonas | genus |
| 169973 | uncultured Massilia sp. | species |
| 69278 | Aquamicrobium | genus |
| 1817 | Nocardia | genus |
| 53458 | Janibacter limosus | species |
| 460257 | Aeromicrobium ponti | species |
| 1827 | Rhodococcus | genus |
| 311234 | Cellulomonas terrae | species |
| 59732 | Chryseobacterium | genus |

| | | |
|---|---|---|
| 1707 | Cellulomonas | genus |
| 460257 | Aeromicrobium ponti | species |
| 16 | Methylophilus | genus |
| 92511 | Curtobacterium sp. B20 | species |
| 257003 | Sphingomonas phyllosphaerae | species |
| 460257 | Aeromicrobium ponti | species |
| 454586 | Pedobacter agri | species |
| 255475 | Aurantimonadaceae | family |
| 16 | Methylophilus | genus |
| 16 | Methylophilus | genus |
| 85413 | Bosea | genus |
| 1385 | Bacillales | order |
| 255475 | Aurantimonadaceae | family |
| 2062 | Streptomycetaceae | family |
| 367298 | Phycicoccus | genus |
| 84567 | Pedobacter | genus |
| 34038 | Rahnella aquatilis | species |
| 162491 | Oerskovia | genus |
| 80864 | Comamonadaceae | family |
| 1121270 | evo_1030728.001 | strain |
| 1121270 | evo_1030728.009 | strain |
| 1121270 | evo_1030728.011 | strain |
| 1121270 | evo_1030728.035 | strain |
| 1121270 | evo_1030728.038 | strain |
| 745369 | evo_1035921.007 | strain |
| 745369 | evo_1035921.008 | strain |
| 745369 | evo_1035921.028 | strain |

| | | |
|---|---|---|
| 745369 | evo_1035921.030 | strain |
| 169973 | evo_1139_Y.018 | strain |
| 169973 | evo_1139_Y.035 | strain |
| 1385 | evo_1286_G.010 | strain |
| 1385 | evo_1286_G.034 | strain |

Supplementary Table 7. List of genomes used in the high complexity dataset, sample 1 (test2high1)

| Taxonomy_ID | Name | Rank |
|---|---|---|
| 478009 | Halobacterium salinarum | species |
| 2037 | Actinomycetales | order |
| 1763 | Mycobacterium | genus |
| 1817 | Nocardia | genus |
| 756689 | Nocardia amikacinitolerans | species |
| 1827 | Rhodococcus | genus |
| 908626 | Angustibacter | genus |
| 1707 | Cellulomonas | genus |
| 162491 | Oerskovia | genus |
| 53458 | Janibacter limosus | species |
| 136099 | Knoellia | genus |
| 367298 | Phycicoccus | genus |
| 53355 | Terrabacter | genus |
| 85023 | Microbacteriaceae | family |
| 33877 | Agromyces | genus |
| 293890 | Agromyces subbeticus | species |
| 2034 | Curtobacterium | genus |
| 96492 | Frigoribacterium | genus |
| 110932 | Leifsonia | genus |

| | | |
|---|---|---|
| 33882 | Microbacterium | genus |
| 190323 | Plantibacter | genus |
| 33886 | Rathayibacter | genus |
| 1663 | Arthrobacter | genus |
| 60919 | Sanguibacter | genus |
| 2040 | Aeromicrobium | genus |
| 1839 | Nocardioides | genus |
| 1843 | Nocardioides jensenii | species |
| 42197 | Actinosynnema pretiosum | species |
| 2030 | Kibdelosporangium aridum | species |
| 68170 | Lechevalieria aerocolonigenes | species |
| 40571 | Lentzea albidocapillata | species |
| 1933 | Streptoalloteichus tenebrarius | species |
| 2062 | Streptomycetaceae | family |
| 2063 | Kitasatospora | genus |
| 1883 | Streptomyces | genus |
| 36874 | Porphyromonas cangingivalis | species |
| 393921 | Porphyromonas crevioricanis | species |
| 1122991 | Prevotella shahii | species |
| 1004 | Chitinophaga sancti | species |
| 279824 | Algoriphagus alkaliphilus | species |
| 990 | Cytophaga xylanolytica | species |
| 292407 | Dyadobacter crusticola | species |
| 76595 | Cellulophaga fucicola | species |
| 59732 | Chryseobacterium | genus |
| 172045 | Elizabethkingia miricola | species |
| 237 | Flavobacterium | genus |

| | | |
|---|---|---|
| 49280 | Gelidibacter algens | species |
| 328515 | Nonlabens dokdonensis | species |
| 906888 | Nonlabens ulvanivorans | species |
| 270918 | Salegentibacter mishustinae | species |
| 435906 | Salegentibacter salarius | species |
| 84567 | Pedobacter | genus |
| 454586 | Pedobacter agri | species |
| 188932 | Pedobacter cryoconitis | species |
| 1385 | Bacillales | order |
| 1386 | Bacillus | genus |
| 1121088 | Bacillus coagulans | species |
| 55080 | Brevibacillus | genus |
| 44249 | Paenibacillus | genus |
| 684063 | Paenibacillus algorifonticola | species |
| 1123358 | Tetragenococcus halophilus | species |
| 1122148 | Lactobacillus lindneri | species |
| 1122150 | Lactobacillus nagelii | species |
| 1122151 | Lactobacillus paralimentarius | species |
| 1121128 | Butyricicoccus pullicaecorum | species |
| 1121320 | Clostridium intestinale | species |
| 1121326 | Clostridium magnum | species |
| 1513 | Clostridium tetani | species |
| 42322 | Eubacterium ruminantium | species |
| 1121131 | Butyrivibrio fibrisolvens | species |
| 1121132 | Butyrivibrio hungatei | species |
| 1123012 | Pseudobutyrivibrio xylanivorans | species |
| 1121421 | Desulfotomaculum aeronauticum | species |

| | | |
|---|---|---|
| 1121428 | Desulfotomaculum hydrothermale | species |
| 445973 | Intestinibacter bartlettii | species |
| 1121324 | Peptoclostridium litorale | species |
| 1525 | Moorella thermoacetica | species |
| 1516 | Thermoanaerobacter thermohydrosulfuricus | species |
| 1120997 | Anaerovibrio lipolyticus | species |
| 349095 | Megasphaera paucivorans | species |
| 41275 | Brevundimonas | genus |
| 75 | Caulobacter | genus |
| 255475 | Aurantimonadaceae | family |
| 414371 | Aureimonas | genus |
| 1121026 | Aureimonas altamirensis | species |
| 85413 | Bosea | genus |
| 53254 | Bosea thiooxidans | species |
| 374 | Bradyrhizobium | genus |
| 46913 | Devosia | genus |
| 1121477 | Devosia limi | species |
| 407 | Methylobacterium | genus |
| 1122234 | Methylobacterium komagatae | species |
| 31988 | Aminobacter | genus |
| 561088 | Aquamicrobium aerolatum | species |
| 68287 | Mesorhizobium | genus |
| 245876 | Nitratireductor | genus |
| 106591 | Ensifer | genus |
| 379 | Rhizobium | genus |
| 382 | Sinorhizobium meliloti | species |
| 1080 | Afifella marina | species |

| | | |
|---|---|---|
| 280 | Xanthobacter autotrophicus | species |
| 990712 | Albidovulum xiamenense | species |
| 576117 | Celeribacter halophilus | species |
| 402884 | Cereibacter changlensis | species |
| 561184 | Mameliella alba | species |
| 266 | Paracoccus denitrificans | species |
| 82367 | Paracoccus pantotrophus | species |
| 1061 | Rhodobacter capsulatus | species |
| 121821 | Roseinatronobacter thiooxidans | species |
| 266809 | Thalassobacter stenotrophicus | species |
| 1120923 | Acidocella aminolytica | species |
| 192 | Azospirillum brasilense | species |
| 1082 | Phaeospirillum fulvum | species |
| 361177 | Altererythrobacter | genus |
| 198312 | Porphyrobacter sanguineus | species |
| 165695 | Sphingobium | genus |
| 13687 | Sphingomonas | genus |
| 563996 | Sphingomonas hankookensis | species |
| 1123269 | Sphingomonas sanxanigenens | species |
| 160791 | Sphingomonas wittichii | species |
| 165697 | Sphingopyxis | genus |
| 506 | Alcaligenaceae | family |
| 80864 | Comamonadaceae | family |
| 12916 | Acidovorax | genus |
| 747294 | Pseudorhodoferax | genus |
| 174951 | Ramlibacter | genus |
| 34072 | Variovorax | genus |

| | | |
|---|---|---|
| 75654 | Duganella | genus |
| 29580 | Janthinobacterium | genus |
| 149698 | Massilia | genus |
| 16 | Methylophilus | genus |
| 915 | Nitrosomonas europaea | species |
| 1121029 | Azoarcus communis | species |
| 1121393 | Desulfatibacillum alkenivorans | species |
| 1121405 | Desulfococcus multivorans | species |
| 91360 | Desulforhopalus singaporensis | species |
| 1121439 | Desulfovibrio alkalitolerans | species |
| 876 | Desulfovibrio desulfuricans | species |
| 1123324 | Succinivibrio dextrinosolvens | species |
| 299255 | Ferrimonas marina | species |
| 543 | Enterobacteriaceae | family |
| 1123238 | Atlantibacter subterranea | species |
| 1121872 | Erwinia tracheiphila | species |
| 53335 | Pantoea | genus |
| 469 | Acinetobacter | genus |
| 170623 | Azotobacter beijerinckii | species |
| 354 | Azotobacter vinelandii | species |
| 286 | Pseudomonas | genus |
| 1123015 | Pseudomonas aeruginosa | species |
| 1123022 | Pseudomonas xiamenensis | species |
| 75309 | Rhodanobacter | genus |
| 32033 | Xanthomonadaceae | family |
| 1121014 | Arenimonas donghaensis | species |
| 68 | Lysobacter | genus |

| | | |
|---|---|---|
| 1122185 | Lysobacter concretionis | species |
| 83618 | Pseudoxanthomonas | genus |
| 40323 | Stenotrophomonas | genus |
| 338 | Xanthomonas | genus |
| 2371 | Xylella fastidiosa | species |
| 48467 | Prosthecobacter debontii | species |

Supplementary Table 8. List of genomes used in the high complexity dataset, sample 2 (test2high2)

| Taxonomy_ID | Name | Rank |
|---|---|---|
| 478009 | Halobacterium salinarum | species |
| 2037 | Actinomycetales | order |
| 1763 | Mycobacterium | genus |
| 1817 | Nocardia | genus |
| 756689 | Nocardia amikacinitolerans | species |
| 1827 | Rhodococcus | genus |
| 908626 | Angustibacter | genus |
| 1707 | Cellulomonas | genus |
| 162491 | Oerskovia | genus |
| 136099 | Knoellia | genus |
| 367298 | Phycicoccus | genus |
| 53355 | Terrabacter | genus |
| 85023 | Microbacteriaceae | family |
| 33877 | Agromyces | genus |
| 293890 | Agromyces subbeticus | species |
| 2034 | Curtobacterium | genus |
| 96492 | Frigoribacterium | genus |
| 110932 | Leifsonia | genus |

| | | |
|---|---|---|
| 33882 | Microbacterium | genus |
| 190323 | Plantibacter | genus |
| 33886 | Rathayibacter | genus |
| 1663 | Arthrobacter | genus |
| 60919 | Sanguibacter | genus |
| 2040 | Aeromicrobium | genus |
| 1839 | Nocardioides | genus |
| 1843 | Nocardioides jensenii | species |
| 42197 | Actinosynnema pretiosum | species |
| 2030 | Kibdelosporangium aridum | species |
| 68170 | Lechevalieria aerocolonigenes | species |
| 40571 | Lentzea albidocapillata | species |
| 1933 | Streptoalloteichus tenebrarius | species |
| 2062 | Streptomycetaceae | family |
| 2063 | Kitasatospora | genus |
| 1883 | Streptomyces | genus |
| 36874 | Porphyromonas cangingivalis | species |
| 393921 | Porphyromonas crevioricanis | species |
| 1122991 | Prevotella shahii | species |
| 1004 | Chitinophaga sancti | species |
| 279824 | Algoriphagus alkaliphilus | species |
| 292407 | Dyadobacter crusticola | species |
| 76595 | Cellulophaga fucicola | species |
| 59732 | Chryseobacterium | genus |
| 172045 | Elizabethkingia miricola | species |
| 237 | Flavobacterium | genus |
| 49280 | Gelidibacter algens | species |

| | | |
|---|---|---|
| 328515 | Nonlabens dokdonensis | species |
| 906888 | Nonlabens ulvanivorans | species |
| 270918 | Salegentibacter mishustinae | species |
| 435906 | Salegentibacter salarius | species |
| 84567 | Pedobacter | genus |
| 454586 | Pedobacter agri | species |
| 188932 | Pedobacter cryoconitis | species |
| 1385 | Bacillales | order |
| 1386 | Bacillus | genus |
| 1121088 | Bacillus coagulans | species |
| 55080 | Brevibacillus | genus |
| 44249 | Paenibacillus | genus |
| 684063 | Paenibacillus algorifonticola | species |
| 759851 | Sporosarcina newyorkensis | species |
| 1123358 | Tetragenococcus halophilus | species |
| 1122148 | Lactobacillus lindneri | species |
| 1122150 | Lactobacillus nagelii | species |
| 1122151 | Lactobacillus paralimentarius | species |
| 1121128 | Butyricicoccus pullicaecorum | species |
| 1121264 | Caloranaerobacter azorensis | species |
| 1121320 | Clostridium intestinale | species |
| 1121326 | Clostridium magnum | species |
| 1513 | Clostridium tetani | species |
| 42322 | Eubacterium ruminantium | species |
| 1121131 | Butyrivibrio fibrisolvens | species |
| 1121132 | Butyrivibrio hungatei | species |
| 1123012 | Pseudobutyrivibrio xylanivorans | species |

| | | |
|---|---|---|
| 1121428 | Desulfotomaculum hydrothermale | species |
| 1121324 | Peptoclostridium litorale | species |
| 1525 | Moorella thermoacetica | species |
| 1120997 | Anaerovibrio lipolyticus | species |
| 349095 | Megasphaera paucivorans | species |
| 41275 | Brevundimonas | genus |
| 75 | Caulobacter | genus |
| 255475 | Aurantimonadaceae | family |
| 414371 | Aureimonas | genus |
| 1121026 | Aureimonas altamirensis | species |
| 85413 | Bosea | genus |
| 53254 | Bosea thiooxidans | species |
| 374 | Bradyrhizobium | genus |
| 46913 | Devosia | genus |
| 1121477 | Devosia limi | species |
| 407 | Methylobacterium | genus |
| 31988 | Aminobacter | genus |
| 561088 | Aquamicrobium aerolatum | species |
| 68287 | Mesorhizobium | genus |
| 245876 | Nitratireductor | genus |
| 106591 | Ensifer | genus |
| 379 | Rhizobium | genus |
| 382 | Sinorhizobium meliloti | species |
| 1080 | Afifella marina | species |
| 990712 | Albidovulum xiamenense | species |
| 576117 | Celeribacter halophilus | species |
| 561184 | Mameliella alba | species |

| | | |
|---|---|---|
| 82367 | Paracoccus pantotrophus | species |
| 441209 | Rhodobaca barguzinensis | species |
| 1061 | Rhodobacter capsulatus | species |
| 121821 | Roseinatronobacter thiooxidans | species |
| 266809 | Thalassobacter stenotrophicus | species |
| 1120923 | Acidocella aminolytica | species |
| 192 | Azospirillum brasilense | species |
| 1082 | Phaeospirillum fulvum | species |
| 361177 | Altererythrobacter | genus |
| 198312 | Porphyrobacter sanguineus | species |
| 165695 | Sphingobium | genus |
| 13687 | Sphingomonas | genus |
| 563996 | Sphingomonas hankookensis | species |
| 257003 | Sphingomonas phyllosphaerae | species |
| 1123269 | Sphingomonas sanxanigenens | species |
| 160791 | Sphingomonas wittichii | species |
| 165697 | Sphingopyxis | genus |
| 506 | Alcaligenaceae | family |
| 80864 | Comamonadaceae | family |
| 12916 | Acidovorax | genus |
| 747294 | Pseudorhodoferax | genus |
| 174951 | Ramlibacter | genus |
| 34072 | Variovorax | genus |
| 75654 | Duganella | genus |
| 29580 | Janthinobacterium | genus |
| 149698 | Massilia | genus |
| 16 | Methylophilus | genus |

| | | |
|---|---|---|
| 915 | Nitrosomonas europaea | species |
| 1121029 | Azoarcus communis | species |
| 1121405 | Desulfococcus multivorans | species |
| 91360 | Desulforhopalus singaporensis | species |
| 1121439 | Desulfovibrio alkalitolerans | species |
| 876 | Desulfovibrio desulfuricans | species |
| 1123324 | Succinivibrio dextrinosolvens | species |
| 543 | Enterobacteriaceae | family |
| 1123238 | Atlantibacter subterranea | species |
| 1121872 | Erwinia tracheiphila | species |
| 53335 | Pantoea | genus |
| 34038 | Rahnella aquatilis | species |
| 469 | Acinetobacter | genus |
| 170623 | Azotobacter beijerinckii | species |
| 354 | Azotobacter vinelandii | species |
| 286 | Pseudomonas | genus |
| 1123015 | Pseudomonas aeruginosa | species |
| 75309 | Rhodanobacter | genus |
| 32033 | Xanthomonadaceae | family |
| 1121014 | Arenimonas donghaensis | species |
| 68 | Lysobacter | genus |
| 1122185 | Lysobacter concretionis | species |
| 83618 | Pseudoxanthomonas | genus |
| 40323 | Stenotrophomonas | genus |
| 338 | Xanthomonas | genus |
| 2371 | Xylella fastidiosa | species |
| 35623 | Acholeplasma oculi | species |

| | | |
|---|---|---|
| 171291 | Mycoplasma verecundum | species |
| 48467 | Prosthecobacter debontii | species |
| 48464 | Prosthecobacter fusiformis | species |

Supplementary Table 9. List of genomes used in the high complexity dataset, sample 3 (test2high3 dataset)

| Taxonomy_ID | Name | Rank |
|---|---|---|
| 478009 | Halobacterium salinarum | species |
| 2037 | Actinomycetales | order |
| 1763 | Mycobacterium | genus |
| 1817 | Nocardia | genus |
| 756689 | Nocardia amikacinitolerans | species |
| 1827 | Rhodococcus | genus |
| 908626 | Angustibacter | genus |
| 1707 | Cellulomonas | genus |
| 162491 | Oerskovia | genus |
| 136099 | Knoellia | genus |
| 367298 | Phycicoccus | genus |
| 53355 | Terrabacter | genus |
| 85023 | Microbacteriaceae | family |
| 33877 | Agromyces | genus |
| 293890 | Agromyces subbeticus | species |
| 2034 | Curtobacterium | genus |
| 96492 | Frigoribacterium | genus |
| 110932 | Leifsonia | genus |
| 33882 | Microbacterium | genus |
| 190323 | Plantibacter | genus |
| 33886 | Rathayibacter | genus |

| | | |
|---|---|---|
| 1663 | Arthrobacter | genus |
| 60919 | Sanguibacter | genus |
| 2040 | Aeromicrobium | genus |
| 1839 | Nocardioides | genus |
| 1843 | Nocardioides jensenii | species |
| 42197 | Actinosynnema pretiosum | species |
| 2030 | Kibdelosporangium aridum | species |
| 68170 | Lechevalieria aerocolonigenes | species |
| 40571 | Lentzea albidocapillata | species |
| 1933 | Streptoalloteichus tenebrarius | species |
| 2062 | Streptomycetaceae | family |
| 2063 | Kitasatospora | genus |
| 1883 | Streptomyces | genus |
| 36874 | Porphyromonas cangingivalis | species |
| 28136 | Prevotella oulorum | species |
| 1122991 | Prevotella shahii | species |
| 1004 | Chitinophaga sancti | species |
| 279824 | Algoriphagus alkaliphilus | species |
| 292407 | Dyadobacter crusticola | species |
| 76595 | Cellulophaga fucicola | species |
| 59732 | Chryseobacterium | genus |
| 172045 | Elizabethkingia miricola | species |
| 237 | Flavobacterium | genus |
| 49280 | Gelidibacter algens | species |
| 328515 | Nonlabens dokdonensis | species |
| 906888 | Nonlabens ulvanivorans | species |
| 270918 | Salegentibacter mishustinae | species |

| | | |
|---|---|---|
| 435906 | Salegentibacter salarius | species |
| 84567 | Pedobacter | genus |
| 454586 | Pedobacter agri | species |
| 188932 | Pedobacter cryoconitis | species |
| 1385 | Bacillales | order |
| 1386 | Bacillus | genus |
| 1121088 | Bacillus coagulans | species |
| 55080 | Brevibacillus | genus |
| 44249 | Paenibacillus | genus |
| 684063 | Paenibacillus algorifonticola | species |
| 759851 | Sporosarcina newyorkensis | species |
| 1122148 | Lactobacillus lindneri | species |
| 1122150 | Lactobacillus nagelii | species |
| 1122151 | Lactobacillus paralimentarius | species |
| 1121128 | Butyricicoccus pullicaecorum | species |
| 1121264 | Caloranaerobacter azorensis | species |
| 1121320 | Clostridium intestinale | species |
| 1121326 | Clostridium magnum | species |
| 1513 | Clostridium tetani | species |
| 42322 | Eubacterium ruminantium | species |
| 1121131 | Butyrivibrio fibrisolvens | species |
| 1121132 | Butyrivibrio hungatei | species |
| 1123012 | Pseudobutyrivibrio xylanivorans | species |
| 1121428 | Desulfotomaculum hydrothermale | species |
| 1121324 | Peptoclostridium litorale | species |
| 1525 | Moorella thermoacetica | species |
| 1516 | Thermoanaerobacter thermohydrosulfuricus | species |

| | | |
|---|---|---|
| 1120997 | Anaerovibrio lipolyticus | species |
| 349095 | Megasphaera paucivorans | species |
| 41275 | Brevundimonas | genus |
| 75 | Caulobacter | genus |
| 255475 | Aurantimonadaceae | family |
| 414371 | Aureimonas | genus |
| 1121026 | Aureimonas altamirensis | species |
| 85413 | Bosea | genus |
| 53254 | Bosea thiooxidans | species |
| 374 | Bradyrhizobium | genus |
| 46913 | Devosia | genus |
| 1121477 | Devosia limi | species |
| 407 | Methylobacterium | genus |
| 31988 | Aminobacter | genus |
| 561088 | Aquamicrobium aerolatum | species |
| 68287 | Mesorhizobium | genus |
| 245876 | Nitratireductor | genus |
| 106591 | Ensifer | genus |
| 379 | Rhizobium | genus |
| 382 | Sinorhizobium meliloti | species |
| 1080 | Afifella marina | species |
| 280 | Xanthobacter autotrophicus | species |
| 990712 | Albidovulum xiamenense | species |
| 576117 | Celeribacter halophilus | species |
| 561184 | Mameliella alba | species |
| 441209 | Rhodobaca barguzinensis | species |
| 1061 | Rhodobacter capsulatus | species |

| | | |
|---|---|---|
| 121821 | Roseinatronobacter thiooxidans | species |
| 266809 | Thalassobacter stenotrophicus | species |
| 1120923 | Acidocella aminolytica | species |
| 192 | Azospirillum brasilense | species |
| 1082 | Phaeospirillum fulvum | species |
| 361177 | Altererythrobacter | genus |
| 165695 | Sphingobium | genus |
| 13687 | Sphingomonas | genus |
| 563996 | Sphingomonas hankookensis | species |
| 257003 | Sphingomonas phyllosphaerae | species |
| 1123269 | Sphingomonas sanxanigenens | species |
| 160791 | Sphingomonas wittichii | species |
| 165697 | Sphingopyxis | genus |
| 1123272 | Sphingorhabdus marina | species |
| 506 | Alcaligenaceae | family |
| 80864 | Comamonadaceae | family |
| 12916 | Acidovorax | genus |
| 747294 | Pseudorhodoferax | genus |
| 174951 | Ramlibacter | genus |
| 34072 | Variovorax | genus |
| 75654 | Duganella | genus |
| 29580 | Janthinobacterium | genus |
| 149698 | Massilia | genus |
| 16 | Methylophilus | genus |
| 915 | Nitrosomonas europaea | species |
| 1121029 | Azoarcus communis | species |
| 1121393 | Desulfatibacillum alkenivorans | species |

| | | |
|---|---|---|
| 1121405 | Desulfococcus multivorans | species |
| 91360 | Desulforhopalus singaporensis | species |
| 1121439 | Desulfovibrio alkalitolerans | species |
| 876 | Desulfovibrio desulfuricans | species |
| 1121391 | Desulfacinum infernum | species |
| 1123324 | Succinivibrio dextrinosolvens | species |
| 299255 | Ferrimonas marina | species |
| 543 | Enterobacteriaceae | family |
| 1121872 | Erwinia tracheiphila | species |
| 53335 | Pantoea | genus |
| 1121119 | Brenneria alni | species |
| 34038 | Rahnella aquatilis | species |
| 469 | Acinetobacter | genus |
| 170623 | Azotobacter beijerinckii | species |
| 354 | Azotobacter vinelandii | species |
| 286 | Pseudomonas | genus |
| 1123015 | Pseudomonas aeruginosa | species |
| 92487 | Thiothrix eikelboomii | species |
| 75309 | Rhodanobacter | genus |
| 32033 | Xanthomonadaceae | family |
| 1121014 | Arenimonas donghaensis | species |
| 68 | Lysobacter | genus |
| 1122185 | Lysobacter concretionis | species |
| 83618 | Pseudoxanthomonas | genus |
| 40323 | Stenotrophomonas | genus |
| 338 | Xanthomonas | genus |
| 2371 | Xylella fastidiosa | species |

| | | |
|---|---|---|
| 48467 | *Prosthecobacter debontii* | species |

Supplementary Table 10. Contingency table between FragGeneScan (FG), Prodigal (PD) and geneRFinder (GF) in test dataset (test1)

| | FG x GF | | FG x PD | | FG x PD | |
|---|---|---|---|---|---|---|
| | Positive | Negative | Positive | Negative | Positive | Negative |
| Positive | 36270 | 14129 | 37287 | 1971 | 37264 | 1994 |
| Negative | 2316 | 2265 | 1299 | 14423 | 13135 | 2587 |
| p-value: 0.00000000 | | | | | | |

Supplementary Table 11. Contingency table between FragGeneScan (FG), Prodigal (PD) and geneRFinder (GF) in low dataset (test2low)

| | FG x GF | | FG x PD | | FG x PD | |
|---|---|---|---|---|---|---|
| | Positive | Negative | Positive | Negative | Positive | Negative |
| Positive | 75206 | 163524 | 70644 | 31888 | 99671 | 2861 |
| Negative | 2190 | 14669 | 6752 | 146305 | 139059 | 13998 |
| p-value: 0.00000000 | | | | | | |

Supplementary Table 12. Contingency table between FragGeneScan (FG), Prodigal (PD) and geneRFinder (GF) in medium dataset (test2medium)

| | FG x GF | | FG x PD | | FG x PD | |
|---|---|---|---|---|---|---|
| | Positive | Negative | Positive | Negative | Positive | Negative |
| Positive | 99974 | 226161 | 93883 | 45675 | 135274 | 4284 |
| Negative | 3182 | 18325 | 9273 | 198811 | 190861 | 17223 |
| p-value: 0.00000000 | | | | | | |

Supplementary Table 13. Contingency table between FragGeneScan (FG), Prodigal (PD) and geneRFinder (GF) in high dataset, sample 01 (test2high1)

| | FG x GF | | FG x PD | | FG x PD | |
|---|---|---|---|---|---|---|
| | Positive | Negative | Positive | Negative | Positive | Negative |

| | | | | | | |
|---|---|---|---|---|---|---|
| Positive | 56651 | 130405 | 53743 | 26327 | 77482 | 2588 |
| Negative | 1851 | 11093 | 4759 | 115171 | 109574 | 10356 |
| p-value: 0.00000000 | | | | | | |

Supplementary Table 14. Contingency table between FragGeneScan (FG), Prodigal (PD) and geneRFinder (GF) in high dataset, sample 02 (test2high2)

| | FG x GF | | FG x PD | | FG x PD | |
|---|---|---|---|---|---|---|
| | Positive | Negative | Positive | Negative | Positive | Negative |
| Positive | 56311 | 130694 | 53433 | 25445 | 76377 | 2501 |
| Negative | 1829 | 11166 | 4707 | 116415 | 110628 | 10494 |
| p-value: 0.00000000 | | | | | | |

Supplementary Table 15. Contingency table between FragGeneScan (FG), Prodigal (PD) and geneRFinder (GF) in high dataset, sample 03 (test2high3)

| | FG x GF | | FG x PD | | FG x PD | |
|---|---|---|---|---|---|---|
| | Positive | Negative | Positive | Negative | Positive | Negative |
| Positive | 56336 | 130460 | 53442 | 25688 | 76460 | 2670 |
| Negative | 1905 | 11299 | 4799 | 116071 | 110336 | 10534 |
| p-value: 0.00000000 | | | | | | |

**OSF Guide**
Repository to reproduce the results of this paper
Link: https://osf.io/g4qk5/

### 1. Benchmark of Prediction Genes

In this folder are all the scripts and results generated to produce the benchmark from CAMI datasets.

#### 1.1. fastas

FASTA with sequences belonging to ORFs extracted from CAMI datasets and returned from CD-HIT.

#### 1.2. interpro

This folder contains:
- original_output: the original files outputted by InterproScan
- ipr: script to extract ORF ids which have associated IPR and the corresponding IDs extracted

#### 1.3. model

This folder contains the RF model trained (saved as a caret object - in R).

#### 1.4. script

This folder contains 5 scripts to reproduce paper experiments with CAMI separated by complexity as follows:
- Rscript test_low.R
- Rscript test_medium.R
- Rscript test_highS01.R
- Rscript test_highS02.R
- Rscript test_highS03.R

Additionally, it contains one script to extract ORFs from genomes. To execute it, it's required to update values for the two following variables inside the code:
- inputFile: Path of FASTA file with the corresponding genome
- outputFile: Path of the output FASTA with sequences of ORFs found in the input file

#### 1.5. tableFeatures

This folder contains one file for each complexity. Each file has all ORFs extracted from original sample along with their features and corresponding labels. ORFs whose IDs were contained in InterproScan output were labeled as positive instance.

#### 1.6. tableSequences

This folder contains auxiliary files that store the genomic content of each ORF extracted.

### 2. GeneRFinder

In this folder are all the implemented scripts and test results of the geneRFinder tool, as well as the results of the Prodigal and FragGeneScan tools.

#### 2.1. modelTrain.R

Script to run train and make a new model.

#### 2.2. PaperPrediction.Rproj

Gene prediction project.

#### 2.3. predictionGenes.R

Script to predict genes.

#### 2.4. src

In this folder can be found the scripts used to predict genes, make models and do tests:

- dataPreProcessing.R: Script to preprocess data
- extractSequences.R: Script to extract sequences
- functions.R: Script which contains all functions used
- getTableFeatures.R: Script to get a table with features from data
- modelGenerating.R: Script to run train and test scripts
- PaperPrediction.Rproj: Gene prediction project
- prediction.R: Script to predict genes
- readFastaFile.R: Script to read FASTA file.
- statistics: folder containing all tests to calculate the results, including the MCNemar test.

#### 2.5. input

Files to produce test (test1) and training (training2) dataset. This folder contains:

- test: Folder with annotations for 12 organisms (complete genome, FASTA file and table features)
- train: Folder with link to download the annotations for 129 organisms (complete genome, FASTA file and table features)

#### 2.6. output

Folder with results. In this folder can be found:

- cami: Table features and table sequences from FASTAS (section 1.1)
- FragGeneScan: Files from FragGeneScan results
- model: Output of training1 dataset
- predict: Predictions of the tools
- Prodigal: Files from Prodigal results
- test: Output of test1 dataset

### 3. Supplementary Data

This folder containing this document.

**geneRFinder-Benchmark**
Repository to find the geneRFinder benchmark
Link: https://sourceforge.net/projects/generfinder-benchmark/

The GeneRFinder-Benchmark is a comprehensive benchmark data for gene prediction which is based on data of CAMI (Critical Assessment of Metagenome Interpretation) and contains labeled data from gene regions.

The benchmark is made up of 9 datasets. For each one of them is provided:

- List of names, taxonomy ID and taxonomic level of the genomes of the organisms that make up the dataset (genomes.csv)
- Set of ORFs extracted from the respective selected genomes (sequences.fasta)
- Ground truth for each of the extracted ORFs (groundtruth.csv)

For information about download, see the documentation.

**geneRFinder**
Repository to find the geneRFinder tool
Link: https://github.com/railorena/geneRFinder

**Installation**
To install geneRFinder, please running the script:

```
Rscript ./src/config.R
```

**Usage**

```
Rscript ./geneRFinder.R -i [fasta_file_name] -o [output_file_name] -t [thread_number] -s [start_type] -n [intergenic]
```

[fasta_file_name]: input file name
[output_file_name]: output file name
[thread_number]: number of thread
[start_type]:
1 - if start codon is ATG
2 - if start codon is ATG, GTG and TTG

[intergenic]:
1 - output without intergenic sequences
2 - output with intergenic sequences

For example,

```
Rscript ./geneRFinder.R -i ./example/final.contigs.fa -o output -t 7 -s 1 -n 1
```

Please, download the src/model.RData file separately, it is a large file.
